# Protein-Ligand Interaction Graphs: Learning from Ligand-Shaped 3D Interaction Graphs to Improve Binding Affinity Prediction

**DOI:** 10.1101/2022.03.04.483012

**Authors:** Marc A. Moesser, Dominik Klein, Fergus Boyles, Charlotte M. Deane, Andrew Baxter, Garrett M. Morris

## Abstract

Graph Neural Networks (GNNs) have recently gained in popularity, challenging molecular fingerprints or SMILES-based representations as the predominant way to represent molecules for binding affinity prediction. Although simple ligand-based graphs alone are already useful for affinity prediction, better performance on multi-target datasets has been achieved with models that incorporate 3D structural information. Most recent advances utilize complex GNN architectures to capture 3D protein-ligand information by incorporating ligand-interacting protein atoms as additional nodes in the graphs; or by building a second protein-based graph in parallel. This expands the graph considerably while obfuscating the shape of the underlying ligand, diminishing the advantage that GNNs have when encoding molecular structures. There is therefore a need for a simple and elegant molecular graph representation that retains the topology of the ligand while simultaneously encoding 3D protein-ligand interactions.

We present Protein-Ligand Interaction Graphs (PLIGs): a simple way of representing atom-atom contacts of 3D protein-ligand complexes as node features for GNNs. PLIGs featurize an atom node in the molecular graph by describing each atom’s properties as well as all atom-atom contacts made with protein atoms within a distance threshold. The edges of the graph are therefore identical to ligand-based graphs, but the nodes encode the 3D protein-ligand contacts. Since PLIGs are applicable to any GNN architecture, we have benchmarked their performance with six different GNN architectures, and compared them to conventional ligand-based graphs and fingerprint-based multi-layer perceptron (MLP) models using the CASF-2016 benchmark set where we found PLIG-based Graph Attention Networks (GATNet) to be the best performing model (*ρ*=0.84, RMSE=1.22 pK). In summary, we created a novel graph-based representation that incorporates 3D structural information into the node features of ligand-shaped molecular graphs. The PLIG representation is simple, elegant, flexible and easily customizable, opening up many possibilities of incorporating other 2D and 3D properties into the graph.

**Access:** The code and implementation for PLIGs and all models can be found at github.com/MarcMoesser/Protein-Ligand-Interaction-Graphs.

## 1. Introduction

In early stage pre-clinical drug discovery, one of the most important properties of a small molecule drug is its binding affinity for the correct protein target. High binding affinity is crucial for the overall efficacy of a drug while the careful design of multi-target affinity profiles is highly desirable to avoid toxicity and side-effects. In addition, higher binding affinity allows drugs to be administered at lower doses to generate the desired efficacy, which reduces overall toxicity and increases practicality. Computer-aided drug design (CADD) has been firmly established as a powerful technique in the drug discovery pipeline, especially machine-learning-based methods ([1]).

Interest in applying machine learning (ML) to the development of more accurate scoring functions that can predict protein-ligand binding affinity has grown in the last decade. Classical scoring functions use physics-based methods (force fields), linear combinations of (semi-)empirical terms, or knowledge-based potentials. They are used in popular docking software such as AutoDock4 ([2, 3]), AutoDock Vina ([4]) or GOLD ([5]) to score 3D ligand poses. While these classical scoring functions perform well in docking and virtual screening tasks, they struggle with binding affinity prediction and ranking tasks ([6, 7]).

More recently, ML-based scoring functions that can outperform classical scoring functions for binding affinity prediction have emerged. These models employ a diverse set of ML architectures and features, from classical ML techniques such as random forests and gradient boosted trees ([8, 9, 10]) to deep learning (DL) models ([11, 12, 13]). As the most accurate scoring functions use many different ML architectures, it is not yet clear how protein-ligand complexes should be represented as features for model training. Older quantitative structure activity relationship (QSAR) models, mostly used against a single protein target, use only ligand-based features such as ECFP fingerprints ([14]). Recent models built with the intent of creating scoring functions that can generalise across multiple proteins and diverse ligands aim to incorporate 3D-structural information into their feature space. The main goal of including structure-based information is to create models with the right inductive bias that can learn the biophysics of protein-ligand interactions, rather than regurgitate biases in their training sets.

Graph-based neural networks (GNN) have recently emerged as a powerful method for binding affinity prediction ([13, 12]). Although simple ligand-based graphs alone are already useful for affinity prediction ([13]) similar to how purely ligand-based scoring functions use random forests ([9]), higher performance for affinity prediction and virtual screening tasks on multi-target datasets has been achieved in graphs that can incorporate 3D-structural information ([12, 15]). In order to encode 3D information, previous studies have chosen to expand the graph itself, by including nodes corresponding to protein atoms that are close to the ligand ([12, 15]). This expands the graph’s complexity considerably while obfuscating the topology of the ligand, diminishing the advantage that GNNs might have when encoding molecular structures. We present Protein-Ligand Interaction Graphs (PLIG) to solve this issue of structural data incorporation into GNNs. PLIGs can incorporate 3D-interactions directly into the atom nodes of molecular graphs, encoding the intermolecular contacts made by each ligand atom in the 3D protein-ligand complex. This enables graphs to retain the shape of the ligand, by only altering node features. We benchmarked a large variety of modern GNN architectures: Graph Convolutional Neural networks (GCN, [16]), Graph Attention Networks (GATNet, [17]), Graph Isomorphism Networks (GIN, [18]) a combined GAT-GCN network [13], Graph SAGE [19] and Simple Graph Convolutional Networks (SGC, [20]). GATNet performs best when featurized with PLIGs, outperforming other well known scoring functions such as PLEC ([10]), *K*_DEEP_ ([11]) and graph-based SIGN ([12]) when tested on the CASF-2016 benchmark set.

## 2. Material & Methods

### 2.1 Training and test sets

We investigated the ability of PLIGs with different graph-based neural networks (GNN) to predict the protein-ligand binding affinity. For training and testing, we used the PDBbind database ([21, 22]), a curated set of 3D-structures of protein-ligand complexes and their corresponding experimentally determined binding affinity, obtained from the Protein Data Bank (PDB, [23]). In order to utilize the most up-to-date data, we used the PDBBind 2020 General Set and supplemented it with data from the PDBBind 2016 Refined Set for a total of 19451 protein-ligand complexes. Additionally, for evaluation against docked poses rather than crystal poses, we subjected the combined dataset to the pre-processing and docking procedure described in the Methods Section 2.2, resulting in 14981 valid, docked protein-ligand complexes with the corresponding crystal poses. The dataset was randomly split into training (14254 data points) and validation (455 data points) sets. We used the PDBbind 2016 “Core Set”, also referred to as the “CASF-2016 set”, as our test set since it was used as the “scoring power” benchmark in the CASF exercise in 2016 ([24]). Our test dataset excluded 13 protein-ligand complexes unable to go through the docking procedure failing either at the ligand preparation or post-docking quality control stage (for a breakdown of the processing see Methods Section 2.2), resulting in 272 valid data points. The CASF-2016 test set has been widely used in the community as a test set for evaluating scoring function performance ([9, 8, 12]), and therefore allows us to compare our models to previously published scoring function benchmarks.

Each model was trained to predict the inhibition constant K_i_, the dissociation constant K_d_, or the half-maximal inhibitory concentration IC_50_, depending on which was provided by PDBBind for each complex. For the purpose of this study, these values were considered as interchangeable and are henceforth referred to as “the binding constant, K”. This practice of combining two or all three different affinity values (IC_50_, K_i_ and K_d_) has been previously used for similar studies ([9, 8]). For training and performance evaluation, the negative base-10 logarithm of K, commonly denoted as pK was used:

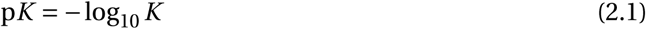

For every model, we evaluated its performance by computing the Pearson correlation coefficient (*ρ*) as well as the root-mean-square error (RMSE in pK units) between the predicted pK value and the experimentally determined pK value for every complex in the test set. Cross validation was performed using a 5-fold split of the training set excluding the validation and test set. Details about cross validation can be found in SI Section 3. Performance on the CASF-2016 benchmark was evaluated for every model by training and testing 10 times using different random seeds on the same dataset split. Then, the *ρ* and RMSE were calculated and reported as the average over 10 runs with the corresponding standard deviation indicating the stability of the model. Model runs were also combined into ensemble models, by averaging each protein-ligand prediction between each of the 10 models in the test set and calculating the *ρ* or RMSE between the average prediction and the true value.

### 2.2 Protein-ligand docking methods

First, ligand files provided by PDBbind were processed using the cheminformatics toolkit RDKit (https://www.rdkit.org/, v2021.03.5). To avoid the starting conformation of the ligand biasing the docking process, new conformers were generated using the ETKDG method ([25]) followed by energy minimization with the Merck Molecular Force Field (MMFF, [26]) as implemented in RDKit. Out of the 19451 original ligand files, some could not be processed by RDKit, and the remaining ligands were subjected to additional filtering criteria (removal of metal containing ligands, removal of molecules with more than 20 rotatable bonds and a molecular weight of more than 1000 Da) resulting in 15317 compounds. This step was performed to filter out non-drug-like ligands. The resulting 15317 ligands were processed into PDBQT-formatted files using Open Babel v3.1.0 ([27]) and the corresponding protein PDB file processed into PDBQT-formatted files using the *prepare_receptor4*.*py* function implemented in MGLTools v1.5.7 ([2]), resulting in one PDBQT file which could not be generated by the software (PDB code 4BPS). The ligand and protein PDBQT files were then docked using Smina ([28]), a fork of AutoDock Vina ([4]) using default parameters except for the following: *exhaustiveness* = 20; *autobox_add* = 8; and *num_modes* = 20. The grid box for each docking run was determined using Smina’s “autobox ligand” feature by passing the original crystal pose of the ligand into Smina and calculating the grid box from its location. For each ligand, 20 diverse poses were generated and the best scoring pose was used for featurization in our models. After docking, the resulting PDBQT ligand files (which lack bond order information) were parsed by RDKit to assign the correct bond orders to the docked pose using the original compound SMILES string. This resulted in the exclusion of 336 ligands and a final set of 14981 valid docked protein-ligand complexes. In order to assess the quality of the docked poses, we computed the root-mean-square deviation (RMSD) between the coordinates of each atom in the docked pose and its corresponding atom in the original crystal pose using the symmetry-aware RMSD method implemented in the Open Drug Discovery Toolkit (ODDT, [29]).

### 2.3 Architecture of machine-learning models

We explored a wide variety of graph-based neural networks (GNNs) as well as a multilayer perceptron neural network (MLPNet) in a two-branch setup (Figure 2.1), with either the GNN or MLPNet embedding the ligand and/or protein structure in the first branch and a second branch encoding the protein sequence in a 1D convolutional neural network. Both branches combine into the read-out layers, consisting of three fully-connected layers. This two-branch architecture is adapted from the GraphDTA architecture described by Nguyen *et al*. [13]. All models were implemented using PyTorch v1.9.0 ([30]) and PyTorch Geometric v2.0.0 ([31]). The architecture and implementation of the graph convolutional neural network (GCN, [16]), graph attention network (GAT, [17]), graph isomorphism network (GIN, [18]), and the combined GAT-GCN were directly adopted from [13] with the only changes being the selection of optimal hyperparameters after hyperparameter tuning (see SI Section 1). In addition, we incorporated GraphSAGE ([19]) as well as Simple Graph Convolutional Networks (SGC, [20]) into the GraphDTA architecture using the PyTorch Geometric SAGEConv and SGCConv implementation respectively. The MLPNet model follows a simple feed-forward neural network architecture implemented using PyTorch with the dimension (width) of each layer and the number of fully-connected layers (depth) determined by hyperparameter optimization as described in SI Section 1. The embedding and setup of the protein branch was adopted from GraphDTA ([13]), with only a minor change: reducing the number of convolutional layers from three layers to only a single layer, simplifying the model. The convolutional kernel size and the number of filters are hyper-parameters which were tuned individually for every GNN/MLPNet and protein branch combination (see SI Section 1). The code for all implemented models can be found at GITHUB ADDRESS.

**Figure 2.1:**
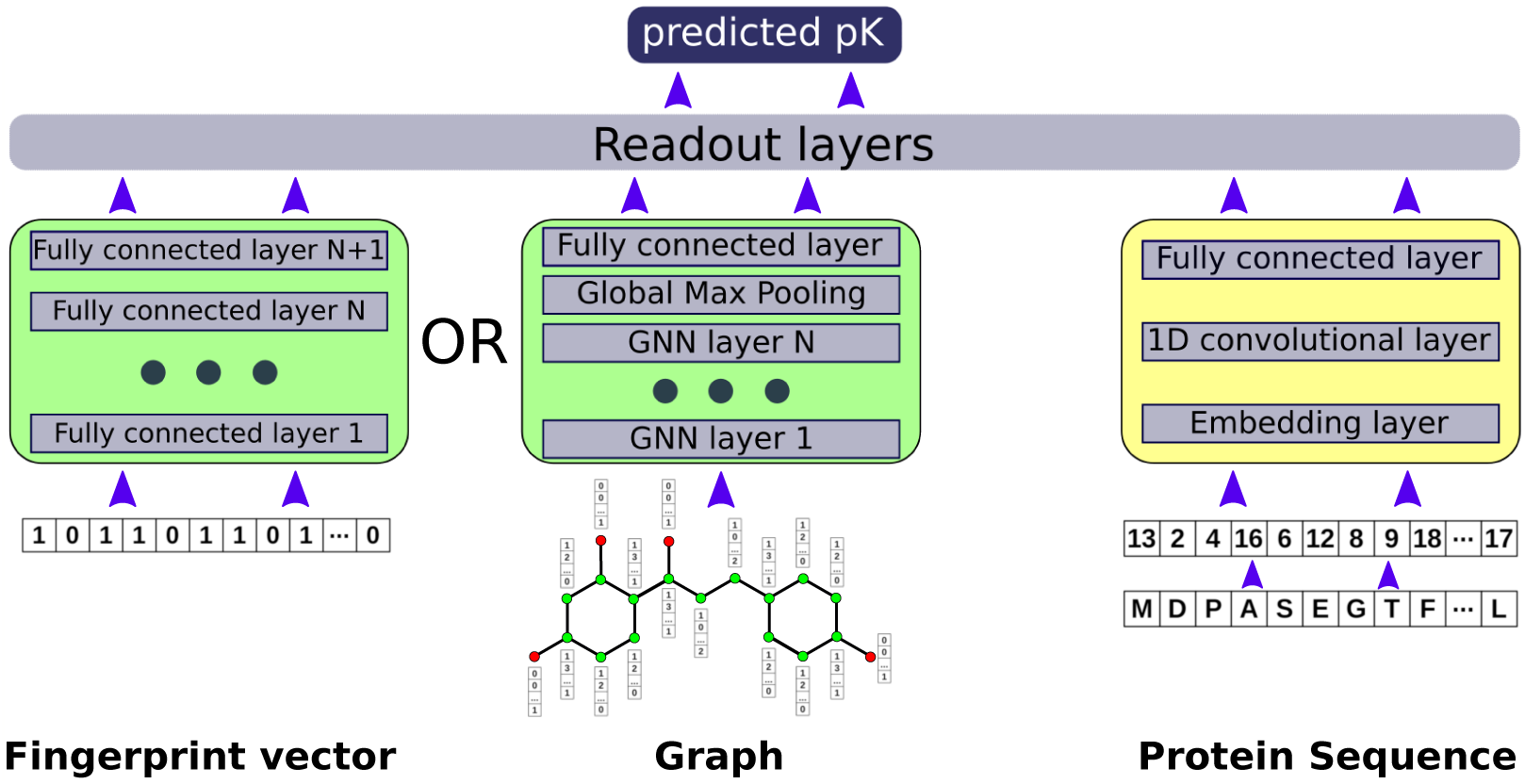
Abstract general architecture of the models with the ligand branch (green) and the sequence branch (yellow). The ligand is either embedded as a fingerprint vector in the MLPNet branch (leftmost branch), or a molecular graph is created for the GNN branch (middle branch). On the right branch, the protein embedding is shown, where a 1D-convolutional layer is fed with the embedded protein amino acid sequence. The outputs from the protein and ligand branches are concatenated and fed into the three fully-connected readout layers and a single pK prediction is made.

### 2.4 Ligand-based graphs

Ligand-based graphs were generated based on the bonds and atoms in a given small molecule. Each atom (node) is represented as a one-hot encoded 40-dimensional feature vector. The feature set used by GraphDTA ([13]) consisted of the following one-hot encoded features: atomic symbol; number of adjacent heavy atoms; number of adjacent hydrogens; implicit valence; and whether the atom is in an aromatic ring. This was expanded by adding additional features: a boolean variable describing whether the atom is in a ring; the one-hot encoded formal charge; and the one-hot encoded hybridization type; and by changing the encoding of the implicit valence to the explicit valence. Since PDBBind ([21, 22]) carefully pre-processes ligands and assigns physiologically relevant atom charges, the inclusion of features such as the explicit valence and the formal charge gives a more complete description of the molecule. All features were calculated using RDKit (https://www.rdkit.org/, v2021.03.5).

### 2.5 Small molecule fingerprints

The molecular fingerprints used for the MLPNet models were calculated using RDKit (https://www.rdkit.org/, v2021.03.5). We investigated the effect of using Morgan fingerprints with the RDKit implementation of Extended-Connectivity Fingerprints and Functional-Class Fingerprints (ECFP and FCFP respectively, [14]) with a radius of 2 and 512 or 1024-bit vectors. They are referred to as “ECFP512” or “ECFP1024” respectively.

### 2.6 Protein-Ligand Interaction Graphs (PLIGs)

The idea of Protein-Ligand Interaction Graphs (or “PLIGs”) was inspired by the work of [8] who introduced Extended Connectivity Interaction Features (ECIF) as high performing features for random forests and gradient boosted tree networks for binding affinity prediction.

PLIGs combine ligand-based graphs with 3D protein-ligand interaction features (Figure 2.2). Initially, a ligand-based graph is generated (see Section (2.4) with only five RDKit-derived atom node features: number of adjacent heavy atoms; number of adjacent hydrogens; explicit valence; aromaticity; and ring membership. Then, all possible protein atom types as defined by Sánchez-Cruz *et al*. [8] in a typical protein (20 proteinogenic amino acids, excluding selenocysteine) were identified based on the following criteria: atom symbol; explicit valence; number of attached heavy atoms; number of attached hydrogens; aromaticity; and ring membership. This results in 22 possible protein atom types (for full list see SI Section 2). To identify the interaction features, the existence of every protein atom in the proximity of a given ligand atom in the 3D-protein-ligand complex is recorded. A 22-dimensional integer vector is created, with each element in the vector corresponding to a unique protein atom type and the integer at that position corresponding to the count of the protein atoms of that type in the vicinity of the ligand atom. Previous work on ECIF ([8]) determined 6 Å to be the optimal proximity threshold, however for this work, we investigated the performance of different thresholds (between 4-8 Åin 1 Å-intervals). Finally, the 5-dimensional ligand-derived atom feature vector is concatenated with the 22-dimensional ligand-protein interaction features, to generate the final 27-dimensional node feature vector. This procedure is repeated for every atom in the ligand to generate the final PLIG.

**Figure 2.2:**
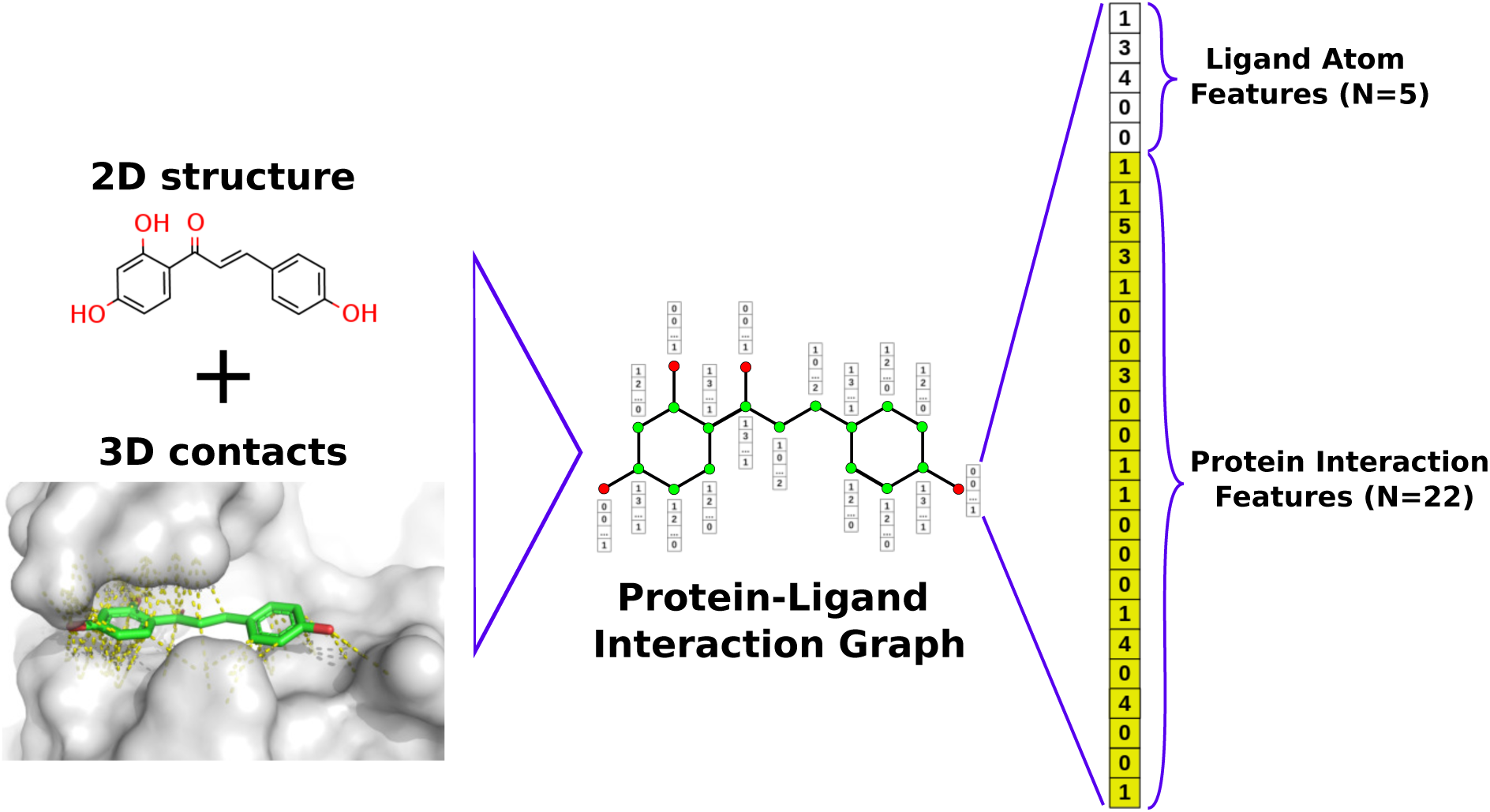
Stylised representation of the construction of a Protein-Ligand Interaction Graph (PLIG) of the PDB structure 6ajv. On the left, the chemical structure of the ligand of 6ajv and 3D-structure of the binding site are shown, with every atom-atom contact in the radius of 4 Å around each ligand atom marked in yellow. The PLIG is then featurized by assigning each atom node to an integer feature vector consisting of the RDKit-derived atom descriptors (5 features: number of adjacent heavy atoms; number of adjacent hydrogens; explicit valence; aromaticity; and ring membership) as well as the number of contacts made with each protein atom type (one for each of the 22 unique atom types in the 20 proteinogenic amino acids) that are within a pre-defined distance threshold to create the final 27-dimensional feature vector for every atom node in the PLIG.

One major disadvantage of the ECIF fingerprints is that the dimensionality of the vector depends on the dataset. An ECIF-featurized model needs to be retrained from scratch every time the model is asked to evaluate a ligand with a new “ECIF atom type” that was not represented in the original dataset. Given the diversity of chemical space, this is highly likely, especially for early stage drug discovery where a diverse set of molecules might be screened or novel scaffolds explored for candidate optimization. PLIGs overcome this limitation by only defining atom types on the protein side, limited to the 20 proteinogenic amino acids (inclusion of more amino acids is also possible if needed), thus eliminating the problems of unknown atom types. Therefore, any new protein-ligand complex can be scored with a pre-trained PLIG model, regardless of the ligand structure. This is especially useful for building prospective models on targets where little structural information is known and docking needed for generating ligand poses, and a diverse set of new ligands needs to be screened. Additionally, the PLIG architecture is able to represent 3D protein-ligand complexes in a ligand-based graph, without needing additional edges and nodes, only changing the node feature vector.

## 3. Results

### 3.1 Quality of Docked Poses

The quality of docked poses was estimated by calculating the symmetry-aware RMSD between all heavy atoms of the original crystal pose and the highest scoring docked pose for each protein-ligand complex using ODDT ([29]). A docked pose is considered to be of high quality if the calculated crystallographic RMSD is 2 Å or less. Overall, 39% of docked poses were high quality. In order to simulate a more realistic docking campaign where lower quality poses can be expected, we used all available docked poses as input for the scoring functions, regardless of pose accuracy. For a discussion of the docking results see SI Section 6.

### 3.2 Model Combinations

For this study, six different GNN and one MLPNet architecture were employed. The GNN models were either featurized using ligand-based graphs or PLIGs. The MLPNet model was featurized with structure-based fingerprints (ECIF [8]) or ligand-based fingerprints (ECFP512, ECFP1024, FCFP512, FCFP1024). All resulting models were trained and tested either in a two-branch architecture with an additional protein sequence encoding branch (see Figure 2.1), or by themselves as a standalone model. This resulted in the 34 models listed in Table 3.1. Each of the models were trained and tested separately on crystallographic and docked poses.

**Table 3.1:** All model architecture and feature combinations. There are 34 different model-feature combinations total. All PLIGS are generated using a proximity threshold of 6 Å.

| Model Architecture | Features | Sequence |
| --- | --- | --- |
| GATNet + PLIG (with sequence) | 3D-based (PLIG) | Yes |
| GATNet + PLIG (no sequence) | 3D-based (PLIG) | No |
| GATNet (with sequence) | ligand-based | Yes |
| GATNet (no sequence) | ligand-based | No |
| GCNNet + PLIG (with sequence) | 3D-based (PLIG) | Yes |
| GCNNet + PLIG (no sequence) | 3D-based (PLIG) | No |
| GCNNet (with sequence) | ligand-based | Yes |
| GCNNet (no sequence) | ligand-based | No |
| GIN + PLIG (with sequence) | 3D-based (PLIG) | Yes |
| GIN + PLIG (no sequence) | 3D-based (PLIG) | No |
| GIN (with sequence) | ligand-based | Yes |
| GIN (no sequence) | ligand-based | No |
| GAT/GCN + PLIG (with sequence) | 3D-based (PLIG) | Yes |
| GAT/GCN + PLIG (no sequence) | 3D-based (PLIG) | No |
| GAT/GCN (with sequence) | ligand-based | Yes |
| GAT/GCN (no sequence) | ligand-based | No |
| SGCNet + PLIG (with sequence) | 3D-based (PLIG) | Yes |
| SGCNet + PLIG (no sequence) | 3D-based (PLIG) | No |
| SGCNet (with sequence) | ligand-based | Yes |
| SGCNet (no sequence) | ligand-based | No |
| SageNet + PLIG (with sequence) | 3D-based (PLIG) | Yes |
| SageNet + PLIG (no sequence) | 3D-based (PLIG) | No |
| SageNet (with sequence) | ligand-based | Yes |
| SageNet (no sequence) | ligand-based | No |
| MLPNet + ECIF (with sequence) | 3D-based (ECIF) | Yes |
| MLPNet + ECIF (no sequence) | 3D-based (ECIF) | No |
| MLPNet + ECFP512 (with sequence) | ligand-based | Yes |
| MLPNet + ECFP512 (no sequence) | ligand-based | No |
| MLPNet + ECFP1024 (with sequence) | ligand-based | Yes |
| MLPNet + ECFP1024 (no sequence) | ligand-based | No |
| MLPNet + FCFP512 (with sequence) | ligand-based | Yes |
| MLPNet + FCFP512 (no sequence) | ligand-based | No |
| MLPNet + FCFP1024 (with sequence) | ligand-based | Yes |
| MLPNet + FCFP1024 (no sequence) | ligand-based | No |

In addition to the models listed in Table 3.1, six different multi-model ensembles were created as described in Section 2.1. The list of created multi-model ensembles is given in Table 3.2. All multi-model ensembles were trained and tested separately on crystallographic and docked poses.

**Table 3.2:**
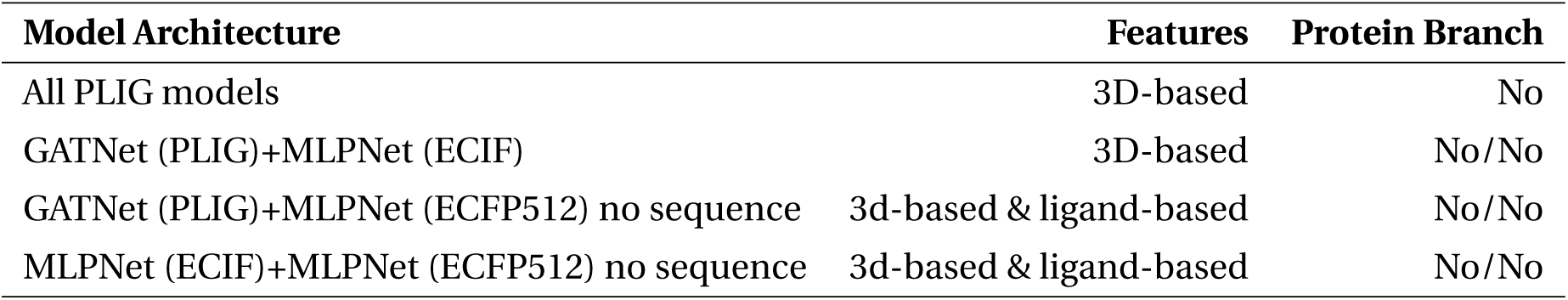
All multi-model ensembles. All models were trained without the sequence embedding. There are 4 multi-model ensembles total. All PLIGS are generated using a proximity threshold of 6 Å.

| Model Architecture | Features | Protein Branch |
| --- | --- | --- |
| All PLIG models | 3D-based | No |
| GATNet (PLIG)+MLPNet (ECIF) | 3D-based | No/No |
| GATNet (PLIG)+MLPNet (ECFP512) no sequence | 3d-based & ligand-based | No/No |
| MLPNet (ECIF)+MLPNet (ECFP512) no sequence | 3d-based & ligand-based | No/No |

### 3.3 Ensemble Model Performance

Since the model predictions are somewhat stochastic, model performance against the withheld test set (CASF-2016) for all trained models was evaluated: (1) using the average and standard deviation (SD) of the Pearson correlation coefficient (*ρ*) as well as the root-mean-square error (RMSE in pK units) over 10 runs; and (2) as the ensemble model over all 10 models by averaging the predictions for each protein-ligand complex in the test set between the 10 models. Model stability between the 10 training runs was high for all models, with a maximum *ρ* standard deviation of 0.023 for the model with highest variability (Ligand-based SGCNet without the protein branch, SI Section 4) and a maximum RMSE standard deviation of 0.08 pK (SGCNet PLIG with and without the protein branch, trained and tested on docked poses). Performance increased for all models when using the ensemble predictions in comparison to the averaged *ρ* score (SI Section 4; Main text Figure 3.1) and therefore all further results are reported for the ensemble models.

**Figure 3.1:**
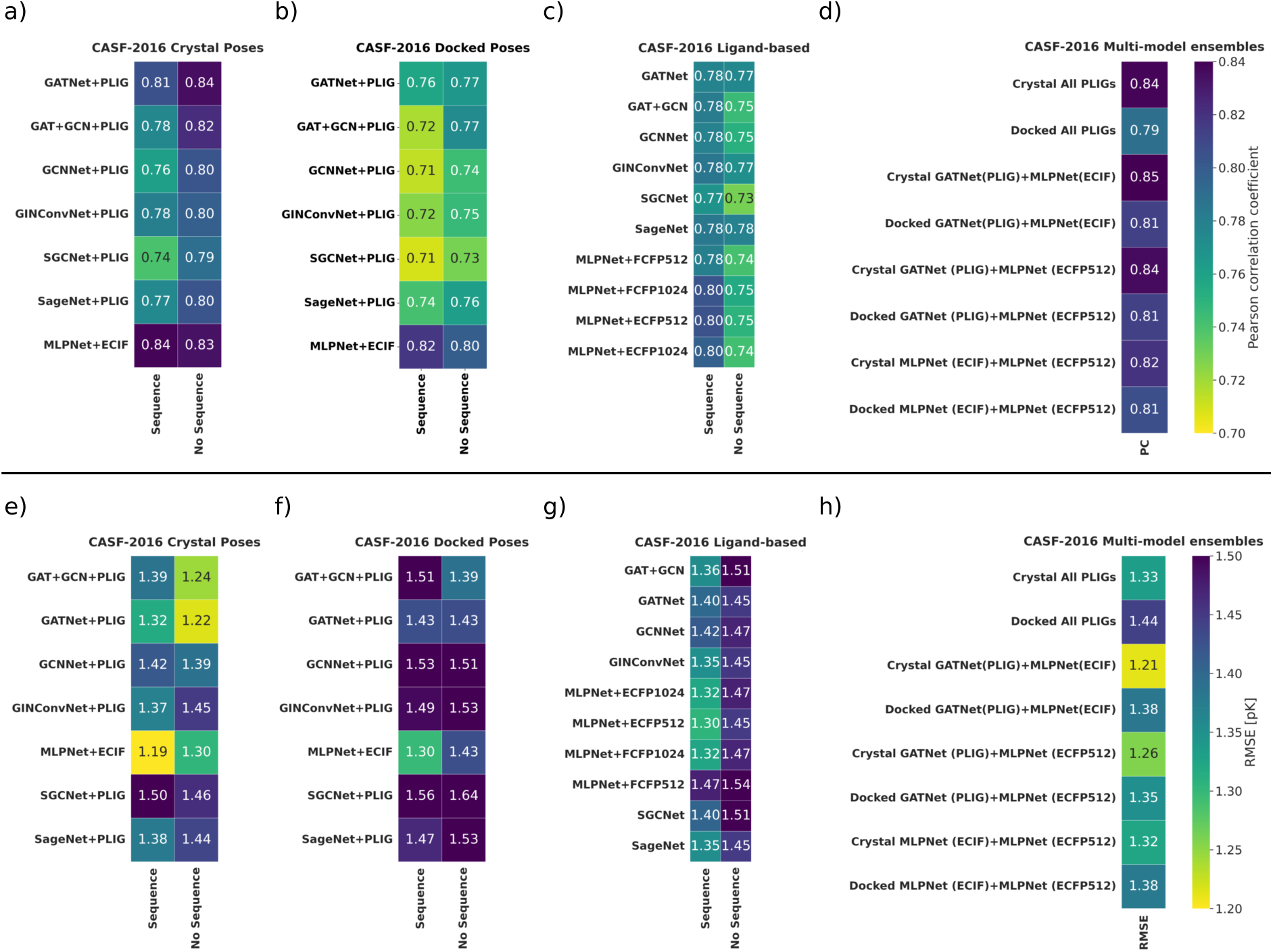
The reported Pearson correlation coefficient (*ρ*) and root-mean-square error (RMSE) are calculated as the ensemble between 10 train and test model iterations by averaging the point predictions for every data point in the test set and calculating *ρ* and RMSEs between the average prediction and the true value. All PLIGs are generated using a proximity threshold of 6 Å. (**a-d**) Pearson correlation coefficients for all model and feature combinations. (**e-h**) RMSE for all model and feature combinations. For both, *ρ* and RMSE results, *“Sequence”* and *“No Sequence”* denote the presence or absence of the protein sequence-encoding branch in the model architecture, respectively. (a and e) Performance of the structure-based GNN and MLPNet models when trained and tested on crystal structures. (b and f) Performance of the structure-based GNN and MLPNet models when trained and tested on docked poses. (c and g) Performance of all ligand-based GNN and MLPNet models. The ECFP and FCFP fingerprint radius was 2 in all cases. (d and h) Performance of the multi-model ensembles for docked and crystal poses. The “All PLIGs” model is an ensemble of all PLIG GNN models. PLIG, ECIF and ECFP models were used without the sequence embedding for all ensembles. The best performing single model, the GATNet PLIG without protein sequence embedding (*ρ*=0.84, RMSE=1.24 pK), was not improved further through multi-model ensembles.

### 3.4 Model Performance on Crystal Poses

Performance of the PLIG and ECIF-based models when trained and tested on crystal poses is shown in Figure 3.1, with the GATNet PLIG model without sequence embedding (*ρ*=0.84, RMSE=1.22 pK) and the MLPNet ECIF model including sequence embedding (*ρ*=0.84, RMSE=1.19 pK) performing best overall in predicting binding affinity. Although performance is strong across all GNN architectures, the GATNet PLIG model outperforms all other graphs. For comparison, [8] report a *ρ* of 0.857 (ECIF without ligand features) and 0.866 (ECIF with additional ligand features) on the CASF-2016 benchmark in a gradient boosted tree model and a previous release of PDBbind (version 2016) as the training data. The performance of ECIF fingerprints is therefore not tied to the original random forest and gradient boosted tree models and can be efficiently used with deep learning as well as in classical machine learning models. We therefore show that (1) PLIGs are a high-performing graph representation of protein-ligand complexes able to elevate GNN-based models to compete with some of the best protein-ligand binding affinity prediction methods such as ECIF ([8]), *K*_DEEP_ ([11]) or AEScore ([32]); and, (2) ECIF are effective in a simple feed forward neural network (MLPNet).

For all PLIG models, the inclusion of the protein sequence branch in the model decreases performance, while it does not significantly affect the performance of the ECIF fingerprint model (Figure 3.1). Since PLIGs already encode the structure of the protein (at least the part near the ligand), inclusion of 1D-protein sequence information might be introducing confounding information or noise, hindering PLIG model performance. In addition, protein-branch containing models are more unstable during cross validation, reaching maximum performance quickly, after a small number of epochs, therefore raising concerns of overfitting (SI Section 3). As a result, we recommend using the single-branch GNN or MLPNet implementations of our models without the protein sequence branch in future studies, with the GATNet PLIG implementation performing best.

### 3.5 Model Performance on Docked Poses

Overall, performance of all models decreased when trained and tested on docked poses in comparison to crystal poses (Figure 3.1 a, b). Like crystal poses, the GATNet PLIG model (no sequence) was the best performing graph (*ρ*=0.77, RMSE=1.43 pK), together with the GAT-GCN PLIG model (no sequence, *ρ*=0.77, RMSE=1.39 pK) when trained and tested on docked poses. However the MLPNet ECIF model (with sequence) performed better (*ρ*=0.82, RMSE=1.30 pK, see Figure 3.1 a, b) only losing performance slightly in comparison to the crystal pose results. As mentioned, models that included the protein sequence embedding were less robust during cross validation (SI Section 3). Therefore, when comparing the MLPNet and GATNet PLIG models without sequence embedding, the difference in *ρ* decreases to 0.03 (MLPNet ECIF no sequence: *ρ*=0.8, GATNet PLIG no sequence: *ρ*=0.77) and the difference in RMSE disappears (RMSE=1.43 pK for both models). The models were also trained on crystal structures and tested on docked poses (SI Figure 4.12) however no significant difference to models trained and tested on docked poses was observed. Overall, the drop in performance from crystal to docked poses for structure-based methods is to be expected, as docked poses tend to diverge from original crystal structures, and contacts identified by those methods might not reflect the actual contacts present in the crystal structure. Nonetheless, GATNet PLIG (no sequence) models perform best among GNN-based methods and are on a par with the MLPNet ECIF (no sequence) model.

### 3.6 Model Performance with Ligand-based Features

When using purely ligand-derived graphs, no significant difference in performance between the different graph architectures was observed. The ligand-based fingerprint models (MLPNet using ECFP and FCFP fingerprints) performed surprisingly well, especially when combined with the sequence embedding, reaching a *ρ* value of up to 0.8 (Figure 3.1 c, MLPNet FCFP1024, ECFP512 and ECFP1024 models). Overall, purely ligand-based models do not perform as well as the structure-based models trained and tested on crystal structures, but perform comparably to the structure-based models trained and tested on docked poses (Figure 3.1 a-c, e-g), showcasing that the additional noise introduced to the dataset via docking leads to models performing as well as ligand-based models that have no 3D data at all. This same trend was observed by Boyles *et al*. [33] for random forest-based models.

### 3.7 Model Performance of Multi-model Ensembles

Since intra-model ensembles increased the performance of all models, we tested the impact of multimodel ensembles. Overall, no significant improvement over the GATNet PLIG model was observed for any model combination (Figure 3.1 d,h) when trained and tested on crystal poses. However, combining GATNet PLIG models with ligand-based ECFP512 fingerprint features yielded an improvement in comparison to the individual models trained and tested on docked poses (ensemble: *ρ*=0.81, RMSE=1.35 pK vs GATNet PLIG (no sequence): *ρ*=0.77, RMSE=1.43; and ECFP512 (no sequence): *ρ*=0.75, RMSE=1.45 pK). Ensembling ligand-based models with MLPNet ECIF models did not increase performance when trained and tested on docked poses. The addition of ligand-based features has been previously shown to recover most of the lost performance when replacing crystal with docked poses (Random Forest models by Boyles *et al*. [33]). We observe the same effect when using graph neural networks and feed forward neural networks (MLPNet) in an ensemble model framework. Being able to apply model ensembles has the advantage that models do not have to be retrained once the decision has been made to include additional ligand features. Rather, in this case, a single new ligand-based model can be created and combined post-prediction with the structure-based models to rescue performance.

Overall, the performance of multi-model ensembles is fairly similar across models, and small differences do not necessarily correlate to superior or inferior models. The GATNet PLIG models perform best, outperforming all other models on crystal poses and when combined with ligand-based models in a multi-model ensemble regain top performance on docked poses.

### 3.8 Influence of Proximity Threshold on Performance

We further investigated the effect of varying the protein-ligand contact proximity thresholds during PLIG generation on the performance of the GATNet PLIG model. We generated PLIGs using a threshold of 4-8 Å in 1 Å steps. These particular thresholds were chosen since values smaller than 4 Å might exclude some atom-atom contacts and therefore give an incomplete picture of the surrounding atom environment. Additionally, proximity values beyond 8 Å would include protein atoms so far away that the relevance of those protein atoms on the ligand atom environment could be considered to be negligible and might introduce more noise in the featurization. The GATNet model without the sequence embedding was chosen as the model architecture for this experiment and 5-fold cross validation as well as training and testing on the CASF-2016 benchmark performed as described above (Methods Section 2.1).

The performance during cross validation as well as the average performance and standard deviation before ensembling is given in SI Section 5. The ensemble performance between 10 train and test model iterations is shown in Table3.3. The best performance when trained and tested on crystal structures was observed for PLIGs with thresholds of 5 Å and 6 Å (*ρ*=0.84 / 0.84 and RMSE=1.21 / 1.22 for the 5 / 6 Å thresholds, respectively) with no significant advantage of one over the other, which is in line with the results observed by Sánchez-Cruz *et al*. [8] who found 6 Å to be the optimal threshold for ECIF fingerprints in random forest and gradient boosted tree models. Furthermore, performance dropped for all thresholds when training and testing on docked poses (Table 3.3) with no significant difference between different model performances. This observation is in line with our expectations since a majority of docked poses were found to have a RMSD between the crystal and docked pose of larger than 2 Å (Section 3.1). A threshold change of the same magnitude should therefore not alter results drastically, as most poses have an equally large inaccuracy in their pose. Finally, multi-model ensembles between all thresholds does not improve performance for crystal poses and only slightly for docked poses. However, this effect is in line with the results observed by multi-model ensembles of all GNN PLIG models (Figure 3.1) and is not a significant improvement. Overall, we found the 6 Å threshold to perform best between docked and crystal poses and recommend usage of 6 Å PLIGs for further studies.

**Table 3.** 3.3: Performance of the GATNet PLIG model on the CASF-2016 benchmark using different proximity thresholds. Reported Pearson correlation coefficient (*ρ*) and Root mean square error (RMSE) are calculated as the ensemble between 10 train and test model iterations by averaging the point predictions for every data point in the test set and calculating *ρ* values and RMSEs between the average prediction and the true value. Best performance on crystal structures were achieved with PLIGs using the 5 and 6 Å thresholds (marked bold, *ρ*=0.84 / 0.84 and RMSE=1.21 / 1.22 for 5 / 6 Å threshold respectively) with no significant difference in performance between both. Thresholds above 6 Å and below 5 Å experience a small drop in performance. However, the difference in performance disappears when training and testing on docked poses where all models perform comparably and experience an overall decrease in performance in comparison to models trained and tested on crystal structures. Multi-model ensembles between all proximity thresholds does not lead to significantly increased performance.

| Model Architecture | Dataset | Ensemble $\rho$ | Ensemble RMSE [pK] |
| --- | --- | --- | --- |
| GATNet (PLIG) 4 Å | Crystal | 0.81 | 1.32 |
| **GATNet (PLIG) 5 Å** | **Crystal** | **0.84** | **1.21** |
| **GATNet (PLIG) 6 Å** | **Crystal** | **0.84** | **1.22** |
| GATNet (PLIG) 7 Å | Crystal | 0.81 | 1.29 |
| GATNet (PLIG) 8 Å | Crystal | 0.80 | 1.33 |
| GATNet (PLIG) 4 Å | Docked | 0.76 | 1.46 |
| GATNet (PLIG) 5 Å | Docked | 0.76 | 1.44 |
| GATNet (PLIG) 6 Å | Docked | 0.77 | 1.43 |
| GATNet (PLIG) 7 Å | Docked | 0.77 | 1.39 |
| GATNet (PLIG) 8 Å | Docked | 0.76 | 1.43 |
| All proximities ensemble | Crystal | 0.84 | 1.22 |
| All proximities ensemble | Docked | 0.79 | 1.38 |

### 3.9 Model Generalizability

In order to assess the ability of the best performing GATNet PLIG model to generalize between different protein families, protein-ligand pairs in the training dataset were eliminated based on their sequence identity to proteins represented in the CASF-2016 benchmark using five threshold levels between 50 % and 100 % identity. The GATNet PLIG (no sequence) model was trained on the reduced dataset and tested against the full CASF-2016 dataset. As expected, performance decreases with stricter identity thresholds (Figure 3.2) with the largest drop in performance observed between the original full dataset and the elimination of 100% identical proteins from the training set (meaning that only proteins in the training set with identical sequence to proteins in the test set are removed). This observation is in line with similar experiments such as reported by Boyles *et al*. [9], where model performance decreased regardless of random forest model featurization when eliminating test set-similar protein structures from the training set. However, part of the decrease in performance could be due to the reduced dataset size when eliminating data points in the training set (SI Table 7.1). Nonetheless, as CASF-2016 is deliberately chosen to be representative of the proteins present in the PDBbind refined set, eliminating this bias through techniques such as the sequence identity elimination described herein, a fairer evaluation of generalizability would be achieved and should therefore be utilized in future evaluations of other protein-ligand affinity scoring functions.

**Figure 3.2:**
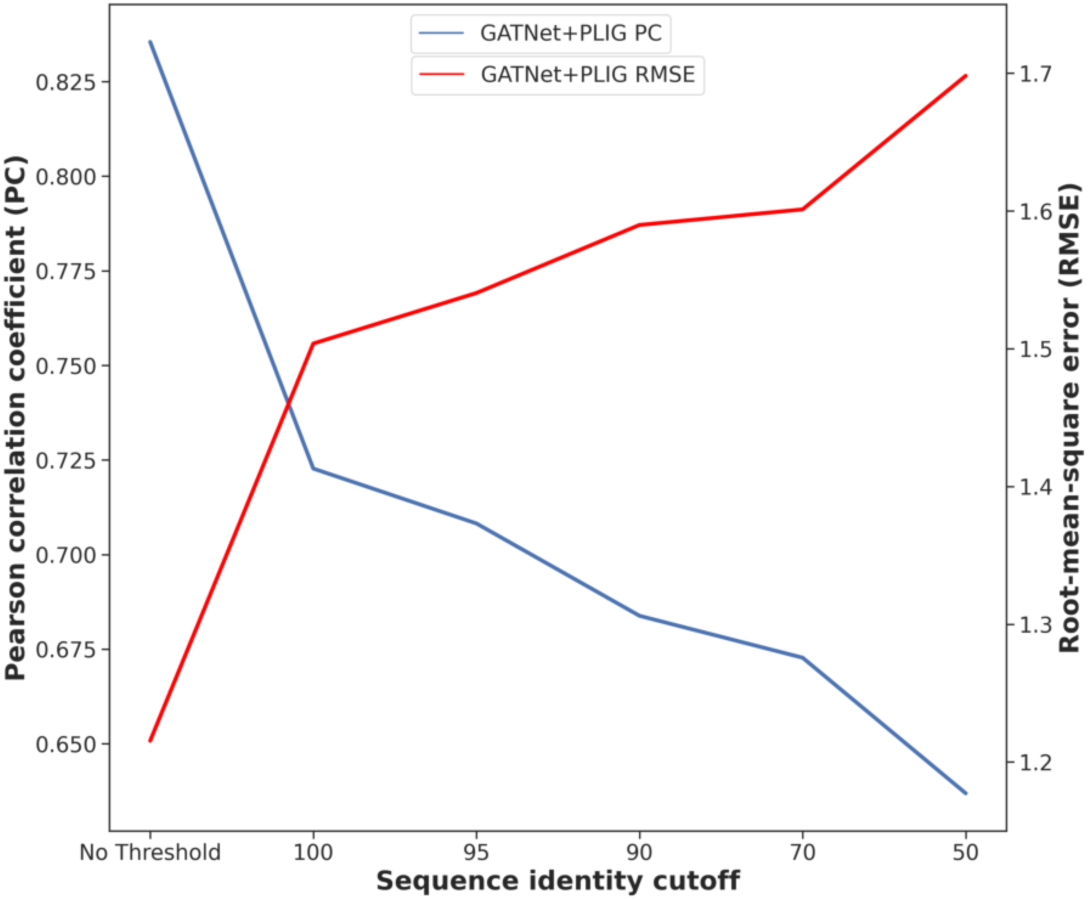
Pearson correlation coefficient (*ρ*) and Root-mean-square error (RMSE) of predicted versus experimental binding affinity for the GATNet PLIG model (no sequence, ensemble over 10 runs) when trained and tested on crystal poses. Protein-ligand complexes in the training set with a sequence identity at or above the cut-off value to proteins in the CASF-2016 test set were excluded resulting in a smaller dataset at every step.

## 4. Discussion

The recent interest in GNN-based scoring functions for protein-ligand binding affinity prediction has led to an arms race in complexity. New models that try to incorporate structural features into GNNs add protein nodes to the graph ([12, 15]) which changes the shape of the graph and introduces unnecessary complexity of the model. In this study, we present Protein-Ligand Interaction Graphs (PLIGs), which are simple molecular graphs that retain the chemical topology of the ligand while also encoding all 3D protein-ligand interactions by incorporating proximity-based contacts into each atom’s node features. Despite their simplicity, PLIGs perform among the top scoring functions when tested against the CASF-2016 benchmark. Using the novel PLIG-based featurization, we conducted a comprehensive analysis of six different GNN architectures and found Graph Attention Networks (GATNet) to be the best for predicting binding affinity using PLIGs. This might be due to the self-attention mechanism of GATs, which allow the weighting factor of each node to be implicitly defined in relation to its neighboring nodes, instead of being explicitly defined or learned. Since the protein contacts made by each atom node, as well as the nodes themselves, are more or less important based on their surrounding neighborhoods, the ability of GATs to elicit contextually-relevant importance of different neighbors could contribute to its increased performance. In addition, simplicity in model design seems to be desirable for PLIG models, since the hybrid branch architecture that combines the protein sequence and the GNN using PLIG performs worse than models with just PLIG-based GNNs.

GATNet PLIGs (*ρ*=0.84, RMSE=1.22) perform comparably to some of the best current scoring functions against the CASF-2016 benchmark such as ECIF fingerprints (*ρ*=0.867, [8]). In addition, PLIGs outperform other well-established structure-based models such as PLEC (*ρ*=0.817, [10]), OnionNet (*ρ*=0.816, [34]) and *K*_DEEP_ (*ρ*=0.82, [11]) as well as other recent GNN-based scoring functions such as SIGN (*ρ*=0.797, [12]).

Finally, we have shown that model ensembles can be a powerful tool in overcoming the performance drop of binding affinity scoring functions when going from training on crystal to training on docked poses. Rather than creating a hybrid model by adding ligand-based features such as ECFP fingerprints directly into the feature space of a structure-based model (as described by Boyles *et al*. [9]), a simple ensemble of a structure-based model and a ligand-based fingerprint model can recover or even increase performance to levels higher than the individual models by themselves. However, ensembling different structure-based methods does not seem to improve performance significantly, at least for the GNN PLIG and MLPNet ECIF models described here. Additional investigation would be required to test if a more diverse set of model ensembles could improve performance further.

This study shows that simply including proximity-based protein-ligand contacts into the atomic nodes of molecular graphs of the ligand boosts performance of graph neural networks when predicting protein-ligand binding affinity. As a result, it opens up a large space for exploration of which other node features might be included in molecular graphs. Rather than increasing the size of the graph, adding nodes or increasing the complexity, this method of incorporating 3D-structural information into ligand graphs is simple, powerful and compatible with a large variety of GNN architectures. Furthermore, the PLIG architecture can serve as a guide to enable researchers to investigate which other kinds of protein-ligand complex representations might be valuable in GNNs by following the steps laid out to create PLIGs, replacing the general contact information with other other kinds of contact information, biophysical quantities or representations of 3D-protein-ligand complex information. Lastly, further study into the feature importance and interpretability of PLIGs could enable a deeper understanding of which individual ligand atoms and protein-ligand contacts drive high affinity since PLIG atom node features are fully interpretable.

## Acknowledgements

We thank the EPSRC University of Oxford Mathematics, Physical, and Life Sciences Division (MPLS) Doctoral Training Partnership (DTP) and GlaxoSmithKline for financial support and the Advanced Research Computing (ARC) service of the University of Oxford as well as the Oxford Protein Informatics Group at the Department of Statistics for access to their high performance compute resources.

