## Supplementary Information for "Protein-Ligand Interaction Graphs: Learning from Ligand-Shaped 3D Interaction Graphs to Improve Binding Affinity Prediction"

**Access:** code implementation for PLIGs can be found at https://github.com/MarcMoesser/Protein-Ligand-Interaction-Graphs

**6 Docking results** **46**

**7 Sequence Similarity Threshold Experiments** **47**

**8 Bibliography** **50**

### 1. *Model hyperparameter tuning*

#### 1.1 Hyperparameter optimization setup

The hyperparameters were optimized using Optuna 2.8.0 [1]. Optimization was performed on the PDBbind refined set 2016 [2, 3], with a random train/validation split resulting in a training dataset of 3229 and a validation dataset of 359 compounds. The CASF-2016 test set that was used as a benchmark set was removed from the PDBBind refined set 2016 prior to hyperparameter optimization. Since each specific architecture has it's own hyperparameters, some changes had to be made between models for optimization. Some hyperparameters appear in every model:

- The learning rate was optimized based on a loguniform prior on $[1e-4, 1e-2]$.
- The activation function in the MLP/GNN layers was optimized with categorical values $\{ReLU, LeakyReLU, Sigmoid\}$.
- The dropout rate for the MLP/GNN layers was optimized based on a uniform prior on $[0, 0.9]$.
- The dropout rate for the readout layers was optimized based on a uniform prior on $[0, 0.9]$.
- The number of MLP/GNN layers was optimized with possible values $\{1, ..., 10\}$.

The readout layers were set to a fixed size as they only serve to combine the features extracted in previous layers. On the protein branch side (if included in the model) the following parameters were optimized:

- The number of filters in the 1D-convolutional layer was optimized with possible values $\{2^2, ..., 2^5\}$.
- The size of the filters in the 1D-convolutional layer was optimized with possible values $\{2^2, ..., 2^5\}$.

The idea behind optimizing these hyperparameters is that the protein branch should not be a bottleneck for the model performance and hence some flexibility should be allowed. In the GNN/MLP branch hyperparameters which were optimized are:

- The number of MLP/GNN layers (except for SGCs) was optimized with possible values $\{1, ..., 10\}$ for GNN models and $\{0, ..., 10\}$ for MLP layers. As the N+1$^{st}$ layer is a fully connected layer $N = 0$ is possible for fingerprint-based models.
- The dimension of each MLP (width of layer) / GNN layer (length of node features) as a multiple of the input dimension was optimized with possible values values $\{F^{(0)} \cdot 2^0, F^{(0)} \cdot 2^1, F^{(0)} \cdot 2^2\}$ where $F^{(0)}$ is the input feature dimension, e.g. $F^{(0)} = 40$ for LBGs and $F^{(0)} = 512$ for the ECFP-512 model.

SGCs are restricted to one GNN layer as their idea is to replace $N$ GCN layers by one SGC layer [4]. Model-specific hyperparameters which are optimized are:

- the power of the adjacency matrix $K$ of SGCs with possible values $\{1, ..., 10\}$,

- the N possibly different dimensions of the GAT heads for GATs with possible values $\{1,...,10\}$,
- the N possibly different dimensions of the GAT heads for GAT-GCNs with possible values $\{1,...,10\}$,
- the initial $\epsilon$-value (uniform on $[0,2]$) for GINs and the MLP activation function of the MLP which is part of a GIN layer among $\{ReLU, LeakyReLU, Sigmoid\}$.

Each hyperparameter optimization comprised 1000 trials with values suggested by the TPE (Tree-structured Parzen Estimator) algorithm which is based on Gaussian Mixture Models [5]. The results for each model can be found in the next section.

#### 1.2 Hyperparameter optimization results

PB included corresponds to the combined 2 branch setup where the protein sequence branch is included. PB Excluded corresponds to the GNN or MLPNet model without protein sequence embedding. We did not hyperparameter tune the MLPNet ECIF models independently and instead utilized the MLPNet ECFP 1024 bit model parameters for the MLPNet ECIF models. The MLPNet ECIF implementation was never intended to get the best possible performance (as [6] have already optimized the performance using random forests) but rather serve as a quick comparison to the new PLIG models as they use similar ECIF style atom representations.

| **Hyperparameter** | **PB Included** | **PB Excluded** |
|---|---:|---:|
| Learning rate | 0.0010 | 0.0006 |
| Activation function | Leaky ReLU | Leaky ReLU |
| Number of filters protein convolution | $2^4$ | - |
| Kernel size of protein convolution | $2^3$ | - |
| Dropout rate GNN layers | 0.033 | 0.000 |
| Dropout rate readout layers | 0.028 | 0.065 |
| Number of GNN layers | 4 | 3 |
| Dimension of GNN layer 1 | $40 \cdot 2^2$ | $40 \cdot 2^2$ |
| Dimension of GNN layer 2 | $40 \cdot 2^2$ | $40 \cdot 2^2$ |
| Dimension of GNN layer 3 | $40 \cdot 2^2$ | $40 \cdot 2^2$ |
| Dimension of GNN layer 4 | $40 \cdot 2^2$ | - |
| Pearson correlation on validation set | 0.759 | 0.729 |

**Table 1.1:** Results of hyperparameter tuning of GCN with ligand-based graphs.

| Hyperparameter | PB Included | PB Excluded |
|---|---|---|
| Learning rate | 0.0008 | 0.0006 |
| Activation function | Leaky ReLU | ReLU |
| Number of filters protein convolution | $2^4$ | – |
| Kernel size of protein convolution | $2^3$ | – |
| Dropout rate GNN layers | 0.021 | 0.110 |
| Dropout rate readout layers | 0.021 | 0.080 |
| Dimension of GNN layer 1 | $40 \cdot 2^0$ | $40 \cdot 2^2$ |
| K | 10 | 10 |
| Pearson correlation on validation set | 0.755 | 0.735 |

**Table 1.2:** Results of hyperparameter tuning of SGC with ligand-based graphs.

| Hyperparameter | PB Included | PB Excluded |
|---|---|---|
| Learning rate | 0.0008 | 0.0007 |
| Activation function | Leaky ReLU | Leaky ReLU |
| Number of filters protein convolution | $2^4$ | – |
| Kernel size of protein convolution | $2^4$ | – |
| Dropout rate GNN layers | 0.016 | 0.029 |
| Dropout rate readout layers | 0.013 | 0.013 |
| Number of GNN layers | 2 | 3 |
| Dimension of GNN layer 1 | $40 \cdot 2^2$ | $40 \cdot 2^0$ |
| Dimension of GAT head 1 | 9 | 10 |
| Dimension of GNN layer 2 | $40 \cdot 2^1$ | $40 \cdot 2^2$ |
| Dimension of GAT head 2 | 1 | 5 |
| Dimension of GNN layer 3 | – | $40 \cdot 2^2$ |
| Dimension of GAT head 3 | – | 6 |
| Pearson correlation on validation set | 0.753 | 0.745 |

**Table 1.3:** Results of hyperparameter tuning of GAT with ligand-based graphs.

| **Hyperparameter** | **PB Included** | **PB Excluded** |
|---|---:|---:|
| Learning rate | 0.0003 | 0.0009 |
| Activation function | Leaky ReLU | ReLU |
| Number of filters protein convolution | $2^4$ | – |
| Kernel size of protein convolution | $2^5$ | – |
| Dropout rate GNN layers | 0.000 | 0.000 |
| Dropout rate readout layers | 0.001 | 0.090 |
| Number of GNN layers | 2 | 1 |
| Dimension of GNN layer 1 | $40 \cdot 2^1$ | $40 \cdot 2^1$ |
| Dimension of GAT head 1 | 5 | 2 |
| Dimension of GNN layer 1 | $40 \cdot 2^1$ | – |
| Dimension of GAT head 1 | 10 | – |
| Pearson correlation on validation set | 0.761 | 0.732 |

**Table 1.4:** Results of hyperparameter tuning of GAT-GCN with ligand-based graphs.

| **Hyperparameter** | **PB Included** | **PB Excluded** |
|---|---:|---:|
| Learning rate | 0.0005 | 0.0003 |
| Activation function | ReLU | ReLU |
| Number of filters protein convolution | $2^4$ | – |
| Kernel size of protein convolution | $2^3$ | – |
| Dropout rate GNN layers | 0.001 | 0.017 |
| Dropout rate readout layers | 0.100 | 0.050 |
| Number of GNN layers | 2 | 3 |
| Dimension of GNN layer 1 | $40 \cdot 2^1$ | $40 \cdot 2^2$ |
| Dimension of GNN layer 2 | $40 \cdot 2^2$ | $40 \cdot 2^1$ |
| Dimension of GNN layer 3 | – | $40 \cdot 2^1$ |
| Initial $\epsilon$ | 0.369 | 0.070 |
| GIN MLP activation function | Leaky ReLU | Leaky ReLU |
| Pearson correlation on validation set | 0.754 | 0.721 |

**Table 1.5:** Results of hyperparameter tuning of GIN with ligand-based graphs.

| Hyperparameter | PB Included | PB Excluded |
|---|---|---|
| Learning rate | 0.0003 | 0.0010 |
| Activation function | Leaky ReLU | Leaky ReLU |
| Number of filters protein convolution | $2^5$ | – |
| Kernel size of protein convolution | $2^5$ | – |
| Dropout rate GNN layers | 0.000 | 0.019 |
| Dropout rate readout layers | 0.016 | 0.147 |
| Number of GNN layers | 6 | 2 |
| Dimension of GNN layer 1 | $40 \cdot 2^1$ | $40 \cdot 2^2$ |
| Dimension of GNN layer 2 | $40 \cdot 2^0$ | $40 \cdot 2^0$ |
| Dimension of GNN layer 3 | $40 \cdot 2^1$ | – |
| Dimension of GNN layer 4 | $40 \cdot 2^0$ | – |
| Dimension of GNN layer 5 | $40 \cdot 2^2$ | – |
| Dimension of GNN layer 6 | $40 \cdot 2^2$ | – |
| Pearson correlation on validation set | 0.755 | 0.734 |

**Table 1.6:** Results of hyperparameter tuning of SAGE with ligand-based graphs.

| Hyperparameter | PB Included | PB Excluded |
|---|---|---|
| Learning rate | 0.0003 | 0.0001 |
| Activation function | ReLU | Leaky ReLU |
| Number of filters protein convolution | $2^3$ | – |
| Kernel size of protein convolution | $2^5$ | – |
| Dropout rate MLP layers | 0.021 | 0.000 |
| Dropout rate readout layers | 0.106 | 0.169 |
| Number of MLP layers | 1 | 6 |
| Dimension of MLP layer 1 | $512 \cdot 2^1$ | $512 \cdot 2^2$ |
| Dimension of MLP layer 2 | – | $512 \cdot 2^0$ |
| Dimension of MLP layer 3 | – | $512 \cdot 2^1$ |
| Dimension of MLP layer 4 | – | $512 \cdot 2^0$ |
| Dimension of MLP layer 5 | – | $512 \cdot 2^2$ |
| Dimension of MLP layer 6 | – | $512 \cdot 2^2$ |
| Pearson correlation on validation set | 0.781 | 0.767 |

**Table 1.7:** Results of hyperparameter tuning of MLP with ECFP-512 fingerprints.

| Hyperparameter | PB Included | PB Excluded |
|---|---|---|
| Learning rate | 0.0001 | 0.0002 |
| Activation function | Leaky ReLU | ReLU |
| Number of filters protein convolution | $2^5$ | – |
| Kernel size of protein convolution | $2^5$ | – |
| Dropout rate MLP layers | 0.019 | 0.099 |
| Dropout rate readout layers | 0.719 | 0.232 |
| Number of MLP layers | 8 | 4 |
| Dimension of MLP layer 1 | $1024 \cdot 2^1$ | $1024 \cdot 2^0$ |
| Dimension of MLP layer 2 | $1024 \cdot 2^0$ | $1024 \cdot 2^0$ |
| Dimension of MLP layer 3 | $1024 \cdot 2^1$ | $1024 \cdot 2^0$ |
| Dimension of MLP layer 4 | $1024 \cdot 2^2$ | $1024 \cdot 2^1$ |
| Dimension of MLP layer 5 | $1024 \cdot 2^1$ | – |
| Dimension of MLP layer 6 | $1024 \cdot 2^1$ | – |
| Dimension of MLP layer 7 | $1024 \cdot 2^1$ | – |
| Dimension of MLP layer 8 | $1024 \cdot 2^2$ | – |
| Pearson correlation on validation set | 0.787 | 0.767 |

**Table 1.8:** Results of hyperparameter tuning of MLP with ECFP-1024 fingerprints.

| Hyperparameter | PB Included | PB Excluded |
|---|---|---|
| Learning rate | 0.0002 | 0.0001 |
| Activation function | ReLU | Leaky ReLU |
| Number of filters protein convolution | $2^4$ | – |
| Kernel size of protein convolution | $2^5$ | – |
| Dropout rate MLP layers | 0.075 | 0.068 |
| Dropout rate readout layers | 0.726 | 0.458 |
| Number of MLP layers | 7 | 8 |
| Dimension of MLP layer 1 | $512 \cdot 2^0$ | $512 \cdot 2^0$ |
| Dimension of MLP layer 2 | $512 \cdot 2^2$ | $512 \cdot 2^1$ |
| Dimension of MLP layer 3 | $512 \cdot 2^1$ | $512 \cdot 2^2$ |
| Dimension of MLP layer 4 | $512 \cdot 2^2$ | $512 \cdot 2^1$ |
| Dimension of MLP layer 5 | $512 \cdot 2^0$ | $512 \cdot 2^0$ |
| Dimension of MLP layer 6 | $512 \cdot 2^2$ | $512 \cdot 2^0$ |
| Dimension of MLP layer 7 | $512 \cdot 2^1$ | $512 \cdot 2^1$ |
| Dimension of MLP layer 8 | – | $512 \cdot 2^1$ |
| Pearson correlation on validation set | 0.775 | 0.761 |

**Table 1.9:** Results of hyperparameter tuning of MLP with FCFP-512 fingerprints.

| Hyperparameter | PB Included | PB Excluded |
|---|---|---|
| Learning rate | 0.0001 | 0.0001 |
| Activation function | Leaky ReLU | ReLU |
| Number of filters protein convolution | $2^5$ | – |
| Kernel size of protein convolution | $2^4$ | – |
| Dropout rate MLP layers | 0.134 | 0.034 |
| Dropout rate readout layers | 0.200[1] | 0.401 |
| Number of MLP layers | 4 | 6 |
| Dimension of MLP layer 1 | $1024 \cdot 2^1$ | $1024 \cdot 2^1$ |
| Dimension of MLP layer 2 | $1024 \cdot 2^1$ | $1024 \cdot 2^1$ |
| Dimension of MLP layer 3 | $1024 \cdot 2^2$ | $1024 \cdot 2^2$ |
| Dimension of MLP layer 4 | $1024 \cdot 2^2$ | $1024 \cdot 2^0$ |
| Dimension of MLP layer 5 | – | $1024 \cdot 2^0$ |
| Dimension of MLP layer 6 | – | $1024 \cdot 2^0$ |
| Pearson correlation on validation set | 0.774 | 0.755 |

**Table 1.10:** Results of hyperparameter tuning of MLP with FCFP-1024 fingerprints.
[1]Result of hyperparameter optimization is 0.842 but since this results in bad performances (especially in combination with early stopping of 5) the value was set to 0.200.

##### Hyperparameter optimization of GNN with PLIG

| Hyperparameter | PB Included | PB Excluded |
|---|---|---|
| Learning rate | 0.0010 | 0.0007 |
| Activation function | Leaky ReLU | Leaky ReLU |
| Number of filters protein convolution | $2^5$ | – |
| Kernel size of protein convolution | $2^3$ | – |
| Dropout rate GNN layers | 0.001 | 0.038 |
| Dropout rate readout layers | 0.017 | 0.058 |
| Number of GNN layers | 5 | 3 |
| Dimension of GNN layer 1 | $27 \cdot 2^1$ | $27 \cdot 2^2$ |
| Dimension of GNN layer 2 | $27 \cdot 2^1$ | $27 \cdot 2^2$ |
| Dimension of GNN layer 3 | $27 \cdot 2^2$ | $27 \cdot 2^0$ |
| Dimension of GNN layer 4 | $27 \cdot 2^2$ | – |
| Dimension of GNN layer 5 | $27 \cdot 2^0$ | – |
| Pearson correlation on validation set | 0.748 | 0.772 |

**Table 1.11:** Results of hyperparameter tuning of GCN with PLIGs.

| Hyperparameter | PB Included | PB Excluded |
|---|---|---|
| Learning rate | 0.0006 | 0.0005 |
| Activation function | Leaky ReLU | ReLU |
| Number of filters protein convolution | $2^3$ | – |
| Kernel size of protein convolution | $2^3$ | – |
| Dropout rate GNN layers | 0.080 | 0.058 |
| Dropout rate readout layers | 0.001 | 0.066 |
| Dimension of GNN layer 1 | $27 \cdot 2^2$ | $27 \cdot 2^2$ |
| K | 2 | 8 |
| Pearson correlation on validation set | 0.761 | 0.765 |

**Table 1.12:** Results of hyperparameter tuning of SGC with PLIGs.

| Hyperparameter | PB Included | PB Excluded |
|---|---|---|
| Learning rate | 0.0001 | 0.0008 |
| Activation function | ReLU | ReLU |
| Number of filters protein convolution | $2^2$ | – |
| Kernel size of protein convolution | $2^2$ | – |
| Dropout rate GNN layers | 0.001 | 0.012 |
| Dropout rate readout layers | 0.034 | 0.020 |
| Number of GNN layers | 3 | 3 |
| Dimension of GNN layer 1 | $27 \cdot 2^1$ | $27 \cdot 2^1$ |
| Dimension of GAT head 1 | 5 | 6 |
| Dimension of GNN layer 2 | $27 \cdot 2^2$ | $27 \cdot 2^2$ |
| Dimension of GAT head 2 | 6 | 5 |
| Dimension of GNN layer 3 | $27 \cdot 2^1$ | $27 \cdot 2^2$ |
| Dimension of GAT head 3 | 9 | 8 |
| Pearson correlation on validation set | 0.786 | 0.787 |

**Table 1.13:** Results of hyperparameter tuning of GAT with PLIGs.

| Hyperparameter | PB Included | PB Excluded |
|---|---|---|
| Learning rate | 0.0008 | 0.0002 |
| Activation function | ReLU | Leaky ReLU |
| Number of filters protein convolution | $2^4$ | – |
| Kernel size of protein convolution | $2^5$ | – |
| Dropout rate GNN layers | 0.004 | 0.000 |
| Dropout rate readout layers | 0.023 | 0.073 |
| Number of GNN layers | 1 | 2 |
| Dimension of GNN layer 1 | $27 \cdot 2^2$ | $27 \cdot 2^2$ |
| Dimension of GAT head 1 | 6 | 8 |
| Dimension of GNN layer 2 | – | $27 \cdot 2^2$ |
| Dimension of GAT head 2 | – | 9 |
| Pearson correlation on validation set | 0.761 | 0.776 |

**Table 1.14:** Results of hyperparameter tuning of GAT-GCN with PLIGs.

| Hyperparameter | PB Included | PB Excluded |
|---|---|---|
| Learning rate | 0.0007 | 0.0012 |
| Activation function | ReLU | Leaky ReLU |
| Number of filters protein convolution | $2^2$ | – |
| Kernel size of protein convolution | $2^5$ | – |
| Dropout rate GNN layers | 0.000 | 0.000 |
| Dropout rate readout layers | 0.014 | 0.307 |
| Number of GNN layers | 1 | 1 |
| Dimension of GNN layer 1 | $27 \cdot 2^2$ | $27 \cdot 2^2$ |
| Initial $\epsilon$ | 1.583 | 1.509 |
| GIN MLP activation function | sigmoid | sigmoid |
| Pearson correlation on validation set | 0.761 | 0.769 |

**Table 1.15:** Results of hyperparameter tuning of GIN with PLIGs.

| Hyperparameter | PB Included | PB Excluded |
|---|---|---|
| Learning rate | 0.0007 | 0.0003 |
| Activation function | ReLU | Leaky ReLU |
| Number of filters protein convolution | $2^5$ | – |
| Kernel size of protein convolution | $2^3$ | – |
| Dropout rate GNN layers | 0.000 | 0.020 |
| Dropout rate readout layers | 0.018 | 0.253 |
| Number of GNN layers | 3 | 5 |
| Dimension of GNN layer 1 | $27 \cdot 2^2$ | $27 \cdot 2^1$ |
| Dimension of GNN layer 2 | $27 \cdot 2^2$ | $27 \cdot 2^1$ |
| Dimension of GNN layer 3 | $27 \cdot 2^2$ | $27 \cdot 2^2$ |
| Dimension of GNN layer 4 | – | $27 \cdot 2^2$ |
| Dimension of GNN layer 5 | – | $27 \cdot 2^2$ |
| Pearson correlation on validation set | 0.768 | 0.762 |

**Table 1.16:** Results of hyperparameter tuning of SAGE with PLIGs.

### 2. *Protein atom types*

The 22 ECIF protein atom types were identified based on the identifiers outlined by [6] from the PDBbind 2020+2016 combined dataset. An atom is defined through the following parameters: atom symbol, explicit valence, number of attached heavy atoms, number of attached hydrogens, aromaticity and ring membership. The possible protein atom types based on the naturally occurring amino acids utilized in PLIG identified based on those parameters is shown in figure 2.1. Out of all possible atom types based on the amino acids only 22 unique atom types are identified. The unique atom types are shown in table 2.1. Those atom types form the dimensionality of the PLIG contact vector of the node features.

| |
|---|
| C;4;2;2;0;0 |
| N;4;1;2;0;0 |
| S;2;2;0;0;0 |
| C;4;3;0;1;1 |
| N;3;1;2;0;0 |
| C;4;3;0;0;0 |
| C;4;1;3;0;0 |
| N;3;2;1;1;1 |
| S;2;1;1;0;0 |
| C;4;3;1;0;1 |
| N;3;2;0;1;1 |
| C;4;3;1;0;0 |
| O;2;1;1;0;0 |
| N;4;2;1;0;0 |
| N;3;2;1;0;0 |
| N;4;1;3;0;0 |
| C;6;3;0;0;0 |
| C;4;2;2;0;1 |
| C;4;2;1;1;1 |
| O;2;1;0;0;0 |
| N;3;3;0;0;1 |
| C;5;3;0;0;0 |

**Table 2.1:** Unique 22 ECIF atom types present in the 20 proteinogenic amino acids in the PDBbind 2020/2016 dataset.

| | Protein Atom | ECIF Atom Type | | Protein Atom | ECIF Atom Type | | Protein Atom | ECIF Atom Type |
|---|---|---|---|---|---|---|---|---|
| **1** | ALA-C | C;4;3;0;0;0 | **63** | GLU-OXT | O;2;1;0;0;0 | **125** | PHE-CZ | C;4;2;1;1;1 |
| **2** | ALA-CA | C;4;3;1;0;0 | **64** | GLY-C | C;4;3;0;0;0 | **126** | PHE-N | N;3;2;1;0;0 |
| **3** | ALA-CB | C;4;1;3;0;0 | **65** | GLY-CA | C;4;2;2;0;0 | **127** | PHE-O | O;2;1;0;0;0 |
| **4** | ALA-N | N;3;2;1;0;0 | **66** | GLY-N | N;3;2;1;0;0 | **128** | PHE-OXT | O;2;1;0;0;0 |
| **5** | ALA-O | O;2;1;0;0;0 | **67** | GLY-O | O;2;1;0;0;0 | **129** | PRO-C | C;4;3;0;0;0 |
| **6** | ALA-OXT | O;2;1;0;0;0 | **68** | GLY-OXT | O;2;1;0;0;0 | **130** | PRO-CA | C;4;3;1;0;1 |
| **7** | ARG-C | C;4;3;0;0;0 | **69** | HIS-C | C;4;3;0;0;0 | **131** | PRO-CB | C;4;2;2;0;1 |
| **8** | ARG-CA | C;4;3;1;0;0 | **70** | HIS-CA | C;4;3;1;0;0 | **132** | PRO-CD | C;4;2;2;0;1 |
| **9** | ARG-CB | C;4;2;2;0;0 | **71** | HIS-CB | C;4;2;2;0;0 | **133** | PRO-CG | C;4;2;2;0;1 |
| **10** | ARG-CD | C;4;2;2;0;0 | **72** | HIS-CD2 | C;4;2;1;1;1 | **134** | PRO-N | N;3;3;0;0;1 |
| **11** | ARG-CG | C;4;2;2;0;0 | **73** | HIS-CE1 | C;4;2;1;1;1 | **135** | PRO-O | O;2;1;0;0;0 |
| **12** | ARG-CZ | C;6;3;0;0;0 | **74** | HIS-CG | C;4;3;0;1;1 | **136** | PRO-OXT | O;2;1;0;0;0 |
| **13** | ARG-N | N;3;2;1;0;0 | **75** | HIS-N | N;3;2;1;0;0 | **137** | SER-C | C;4;3;0;0;0 |
| **14** | ARG-NE | N;4;2;1;0;0 | **76** | HIS-ND1 | N;3;2;0;1;1 | **138** | SER-CA | C;4;3;1;0;0 |
| **15** | ARG-NH1 | N;4;1;2;0;0 | **77** | HIS-NE2 | N;3;2;1;1;1 | **139** | SER-CB | C;4;2;2;0;0 |
| **16** | ARG-NH2 | N;4;1;2;0;0 | **78** | HIS-O | O;2;1;0;0;0 | **140** | SER-N | N;3;2;1;0;0 |
| **17** | ARG-O | O;2;1;0;0;0 | **79** | HIS-OXT | O;2;1;0;0;0 | **141** | SER-O | O;2;1;0;0;0 |
| **18** | ARG-OXT | O;2;1;0;0;0 | **80** | ILE-C | C;4;3;0;0;0 | **142** | SER-OG | O;2;1;1;0;0 |
| **19** | ASN-C | C;4;3;0;0;0 | **81** | ILE-CA | C;4;3;1;0;0 | **143** | SER-OXT | O;2;1;0;0;0 |
| **20** | ASN-CA | C;4;3;1;0;0 | **82** | ILE-CB | C;4;3;1;0;0 | **144** | THR-C | C;4;3;0;0;0 |
| **21** | ASN-CB | C;4;2;2;0;0 | **83** | ILE-CD1 | C;4;1;3;0;0 | **145** | THR-CA | C;4;3;1;0;0 |
| **22** | ASN-CG | C;4;3;0;0;0 | **84** | ILE-CG1 | C;4;2;2;0;0 | **146** | THR-CB | C;4;3;1;0;0 |
| **23** | ASN-N | N;3;2;1;0;0 | **85** | ILE-CG2 | C;4;1;3;0;0 | **147** | THR-CG2 | C;4;1;3;0;0 |
| **24** | ASN-ND2 | N;3;1;2;0;0 | **86** | ILE-N | N;3;2;1;0;0 | **148** | THR-N | N;3;2;1;0;0 |
| **25** | ASN-O | O;2;1;0;0;0 | **87** | ILE-O | O;2;1;0;0;0 | **149** | THR-O | O;2;1;0;0;0 |
| **26** | ASN-OD1 | O;2;1;0;0;0 | **88** | ILE-OXT | O;2;1;0;0;0 | **150** | THR-OG1 | O;2;1;1;0;0 |
| **27** | ASN-OXT | O;2;1;0;0;0 | **89** | LEU-C | C;4;3;0;0;0 | **151** | THR-OXT | O;2;1;0;0;0 |
| **28** | ASP-C | C;4;3;0;0;0 | **90** | LEU-CA | C;4;3;1;0;0 | **152** | TRP-C | C;4;3;0;0;0 |
| **29** | ASP-CA | C;4;3;1;0;0 | **91** | LEU-CB | C;4;2;2;0;0 | **153** | TRP-CA | C;4;3;1;0;0 |
| **30** | ASP-CB | C;4;2;2;0;0 | **92** | LEU-CD1 | C;4;1;3;0;0 | **154** | TRP-CB | C;4;2;2;0;0 |
| **31** | ASP-CG | C;5;3;0;0;0 | **93** | LEU-CD2 | C;4;1;3;0;0 | **155** | TRP-CD1 | C;4;2;1;1;1 |
| **32** | ASP-N | N;3;2;1;0;0 | **94** | LEU-CG | C;4;3;1;0;0 | **156** | TRP-CD2 | C;4;3;0;1;1 |
| **33** | ASP-O | O;2;1;0;0;0 | **95** | LEU-N | N;3;2;1;0;0 | **157** | TRP-CE2 | C;4;3;0;1;1 |
| **34** | ASP-OD1 | O;2;1;0;0;0 | **96** | LEU-O | O;2;1;0;0;0 | **158** | TRP-CE3 | C;4;2;1;1;1 |
| **35** | ASP-OD2 | O;2;1;0;0;0 | **97** | LEU-OXT | O;2;1;0;0;0 | **159** | TRP-CG | C;4;3;0;1;1 |
| **36** | ASP-OXT | O;2;1;0;0;0 | **98** | LYS-C | C;4;3;0;0;0 | **160** | TRP-CH2 | C;4;2;1;1;1 |
| **37** | CYS-C | C;4;3;0;0;0 | **99** | LYS-CA | C;4;3;1;0;0 | **161** | TRP-CZ2 | C;4;2;1;1;1 |
| **38** | CYS-CA | C;4;3;1;0;0 | **100** | LYS-CB | C;4;2;2;0;0 | **162** | TRP-CZ3 | C;4;2;1;1;1 |
| **39** | CYS-CB | C;4;2;2;0;0 | **101** | LYS-CD | C;4;2;2;0;0 | **163** | TRP-N | N;3;2;1;0;0 |
| **40** | CYS-N | N;3;2;1;0;0 | **102** | LYS-CE | C;4;2;2;0;0 | **164** | TRP-NE1 | N;3;2;1;1;1 |
| **41** | CYS-O | O;2;1;0;0;0 | **103** | LYS-CG | C;4;2;2;0;0 | **165** | TRP-O | O;2;1;0;0;0 |
| **42** | CYS-OXT | O;2;1;0;0;0 | **104** | LYS-N | N;3;2;1;0;0 | **166** | TRP-OXT | O;2;1;0;0;0 |
| **43** | CYS-SG | S;2;1;1;0;0 | **105** | LYS-NZ | N;4;1;3;0;0 | **167** | TYR-C | C;4;3;0;0;0 |
| **44** | GLN-C | C;4;3;0;0;0 | **106** | LYS-O | O;2;1;0;0;0 | **168** | TYR-CA | C;4;3;1;0;0 |
| **45** | GLN-CA | C;4;3;1;0;0 | **107** | LYS-OXT | O;2;1;0;0;0 | **169** | TYR-CB | C;4;2;2;0;0 |
| **46** | GLN-CB | C;4;2;2;0;0 | **108** | MET-C | C;4;3;0;0;0 | **170** | TYR-CD1 | C;4;2;1;1;1 |
| **47** | GLN-CD | C;4;3;0;0;0 | **109** | MET-CA | C;4;3;1;0;0 | **171** | TYR-CD2 | C;4;2;1;1;1 |
| **48** | GLN-CG | C;4;2;2;0;0 | **110** | MET-CB | C;4;2;2;0;0 | **172** | TYR-CE1 | C;4;2;1;1;1 |
| **49** | GLN-N | N;3;2;1;0;0 | **111** | MET-CE | C;4;1;3;0;0 | **173** | TYR-CE2 | C;4;2;1;1;1 |
| **50** | GLN-NE2 | N;3;1;2;0;0 | **112** | MET-CG | C;4;2;2;0;0 | **174** | TYR-CG | C;4;3;0;1;1 |
| **51** | GLN-O | O;2;1;0;0;0 | **113** | MET-N | N;3;2;1;0;0 | **175** | TYR-CZ | C;4;3;0;1;1 |
| **52** | GLN-OE1 | O;2;1;0;0;0 | **114** | MET-O | O;2;1;0;0;0 | **176** | TYR-N | N;3;2;1;0;0 |
| **53** | GLN-OXT | O;2;1;0;0;0 | **115** | MET-OXT | O;2;1;0;0;0 | **177** | TYR-O | O;2;1;0;0;0 |
| **54** | GLU-C | C;4;3;0;0;0 | **116** | MET-SD | S;2;2;0;0;0 | **178** | TYR-OH | O;2;1;1;0;0 |
| **55** | GLU-CA | C;4;3;1;0;0 | **117** | PHE-C | C;4;3;0;0;0 | **179** | TYR-OXT | O;2;1;0;0;0 |
| **56** | GLU-CB | C;4;2;2;0;0 | **118** | PHE-CA | C;4;3;1;0;0 | **180** | VAL-C | C;4;3;0;0;0 |
| **57** | GLU-CD | C;5;3;0;0;0 | **119** | PHE-CB | C;4;2;2;0;0 | **181** | VAL-CA | C;4;3;1;0;0 |
| **58** | GLU-CG | C;4;2;2;0;0 | **120** | PHE-CD1 | C;4;2;1;1;1 | **182** | VAL-CB | C;4;3;1;0;0 |
| **59** | GLU-N | N;3;2;1;0;0 | **121** | PHE-CD2 | C;4;2;1;1;1 | **183** | VAL-CG1 | C;4;1;3;0;0 |
| **60** | GLU-O | O;2;1;0;0;0 | **122** | PHE-CE1 | C;4;2;1;1;1 | **184** | VAL-CG2 | C;4;1;3;0;0 |
| **61** | GLU-OE1 | O;2;1;0;0;0 | **123** | PHE-CE2 | C;4;2;1;1;1 | **185** | VAL-N | N;3;2;1;0;0 |
| **62** | GLU-OE2 | O;2;1;0;0;0 | **124** | PHE-CG | C;4;3;0;1;1 | **186** | VAL-O | O;2;1;0;0;0 |
| | | | | | | **187** | VAL-OXT | O;2;1;0;0;0 |

**Figure 2.1:** All heavy atoms in every amino acid in the PDBbind dataset and their corresponding ECIF atom type.

### 3. *Cross validation*

#### 3.1 Cross validation results

5-fold cross validation was performed on the combined PDBBind general 2020 and PDBBind refined 2016 set (dataset details in the main text Section 2.1). The validation and test set (CASF-2016) used in the main study was removed from the cross validation set, to leave the training set of 14254 compounds. Cross validation was done separately for the docked and crystal-based datasets to ensure model stability for both, crystal derived structures and docked poses. This set was split into 5 random folds, using 20 % of the dataset as validation in each fold. Models were run across all 5 folds until model performance converged. The performance of all models during cross validation is recorded for every epoch. The number of epochs to train each model for training and test on the CASF-2016 benchmark was determined as the point where no significant performance increase (difference of less than 0.01 pearson correlation coefficient between epoch $n$ and $n-1$). The determined optimal number of epochs for each model is as follows:

| Model Architecture | PB Included | PB Excluded |
|---|---:|---:|
| GATNet + PLIG | 9 | 16 |
| GCNNet + PLIG | 8 | 18 |
| GIN + PLIG | 6 | 18 |
| GAT/GCN + PLIG | 9 | 19 |
| SGCNet + PLIG | 4 | 14 |
| SageNet + PLIG | 6 | 21 |
| MLPNet + ECIF | 5 | 15 |

**Table 3.1:** Optimal number of epochs for models trained on crystal structures.

| Model Architecture | PB Included | PB Excluded |
|---|---:|---:|
| GATNet + PLIG | 9 | 11 |
| GCNNet + PLIG | 5 | 12 |
| GIN + PLIG | 6 | 17 |
| GAT/GCN + PLIG | 3 | 18 |
| SGCNet + PLIG | 4 | 14 |
| SageNet + PLIG | 8 | 21 |
| MLPNet + ECIF | 5 | 6 |

**Table 3.2:** Optimal number of epochs for models trained on docked poses.

| Model Architecture | PB Included | PB Excluded |
|---|---:|---:|
| GATNet | 4 | 13 |
| GCNNet | 5 | 15 |
| GIN | 7 | 22 |
| GAT/GCN | 7 | 15 |
| SGCNet | 5 | 22 |
| SageNet | 7 | 21 |
| MLPNet + ECFP512 | 5 | 6 |
| MLPNet + ECFP1024 | 7 | 6 |
| MLPNet + FCFP512 | 5 | 8 |
| MLPNet + FCFP1024 | 5 | 4 |

**Table 3.3:** Optimal number of epochs for ligand-based models.

It is noteworthy, that the model architecture that includes the protein branch (PB Included) consistently reaches peak performance significantly faster than the models without protein sequence embedding (all models except the ligand-based ECFP and FCFP fingerprints models using the MLPNet architecture). On average, GNNs using PLIGs and do not include the protein branch take 17 epochs to converge while the same models with the protein branch take only 6 epochs. This could be due to the protein branch models over fitting during training, reaching peak performance quickly, while pure GNN PLIG models need several epochs to slowly learn more meaningful information about the system. The following section includes the performance of all models during cross validation for the reader to inspect.

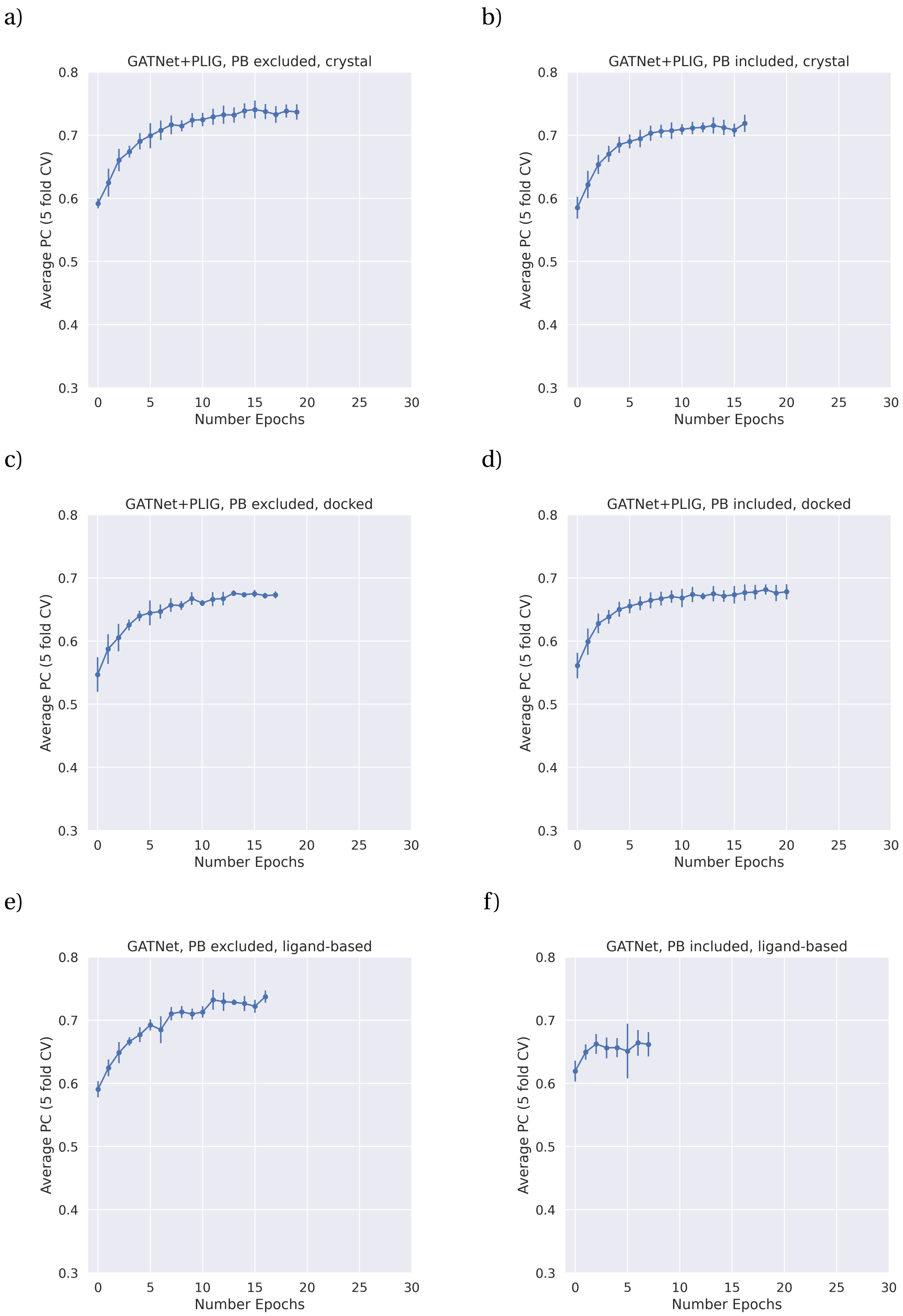

**Figure 3.1:** 5-fold cross validation performance of the GATNet models reported as the average calculated pearson correlation coefficient and it's corresponding standard deviation (error bars) at each epoch. a) PB excluded, crystal structures; b) PB included, crystal structures; c) PB excluded, docked poses; d) PB included, docked poses, e) PB excluded, ligand-based, f) PB included, ligand-based.

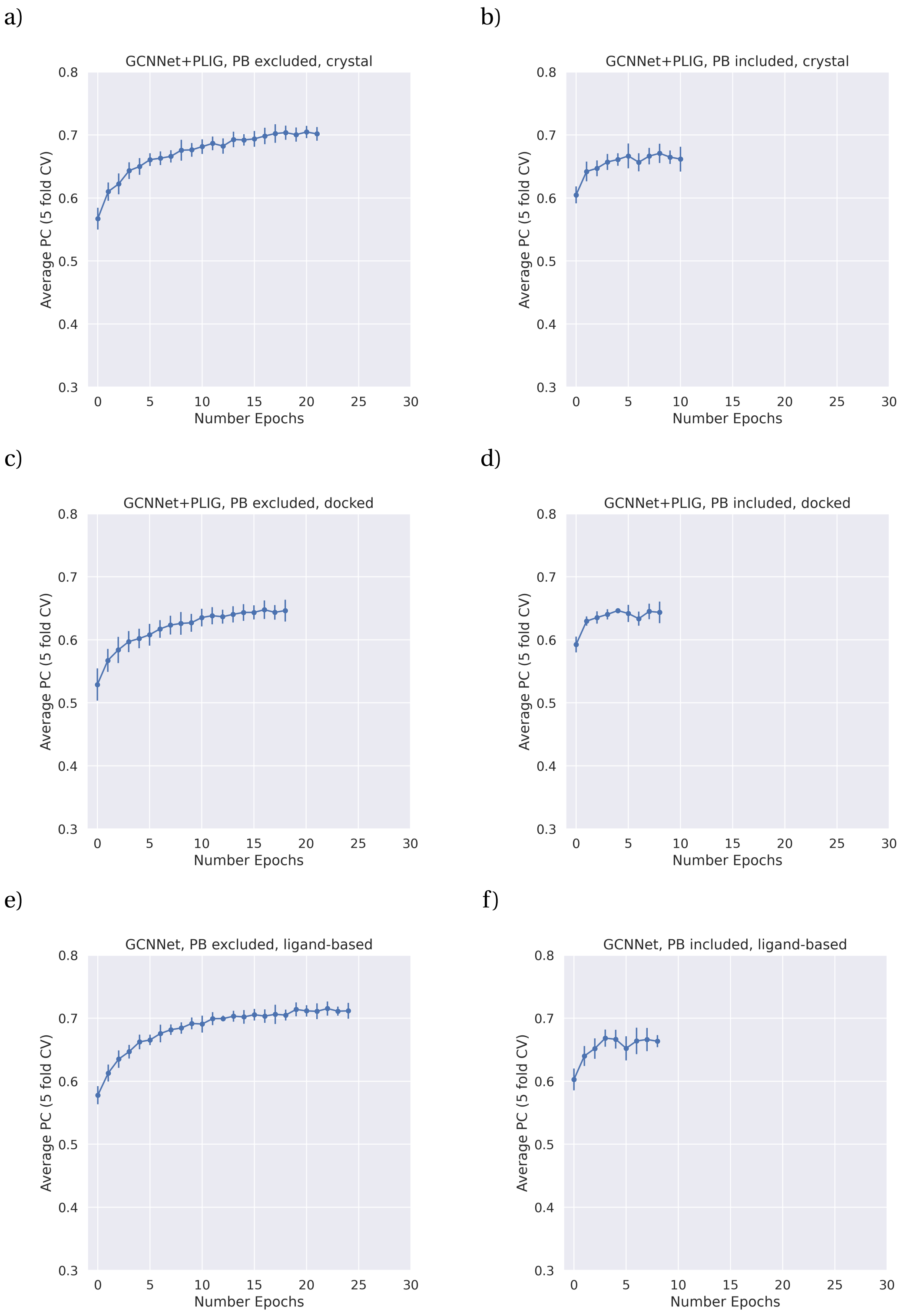

**Figure 3.2:** 5-fold cross validation performance of the GCNNet models reported as the average calculated pearson correlation coefficient and it's corresponding standard deviation (error bars) at each epoch. a) PB excluded, crystal structures; b) PB included, crystal structures; c) PB excluded, docked poses; d) PB included, docked poses, e) PB excluded, ligand-based, f) PB included, ligand-based.

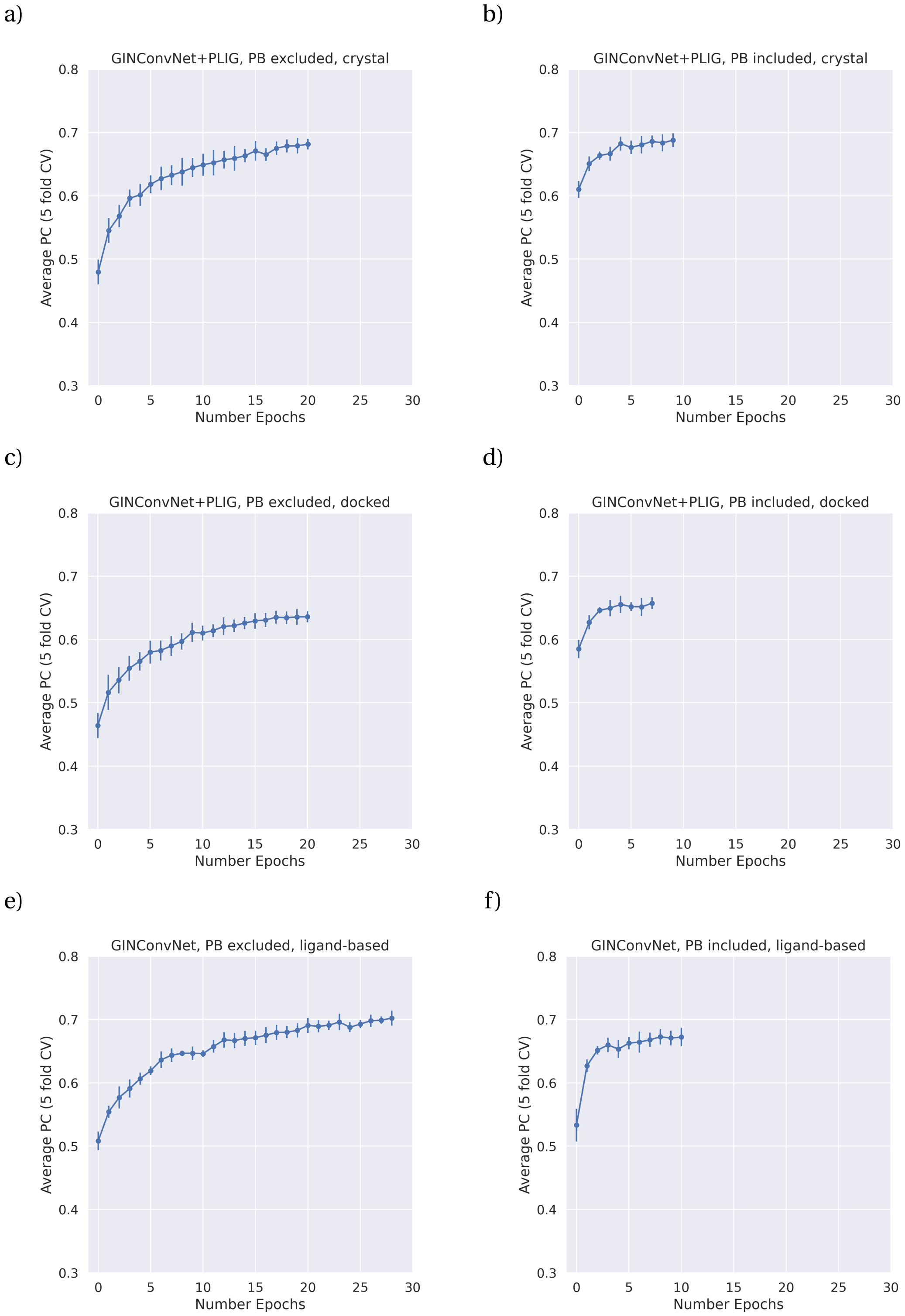

**Figure 3.3:** 5-fold cross validation performance of the GIN models reported as the average calculated pearson correlation coefficient and it's corresponding standard deviation (error bars) at each epoch. a) PB excluded, crystal structures; b) PB included, crystal structures; c) PB excluded, docked poses; d) PB included, docked poses, e) PB excluded, ligand-based, f) PB included, ligand-based.

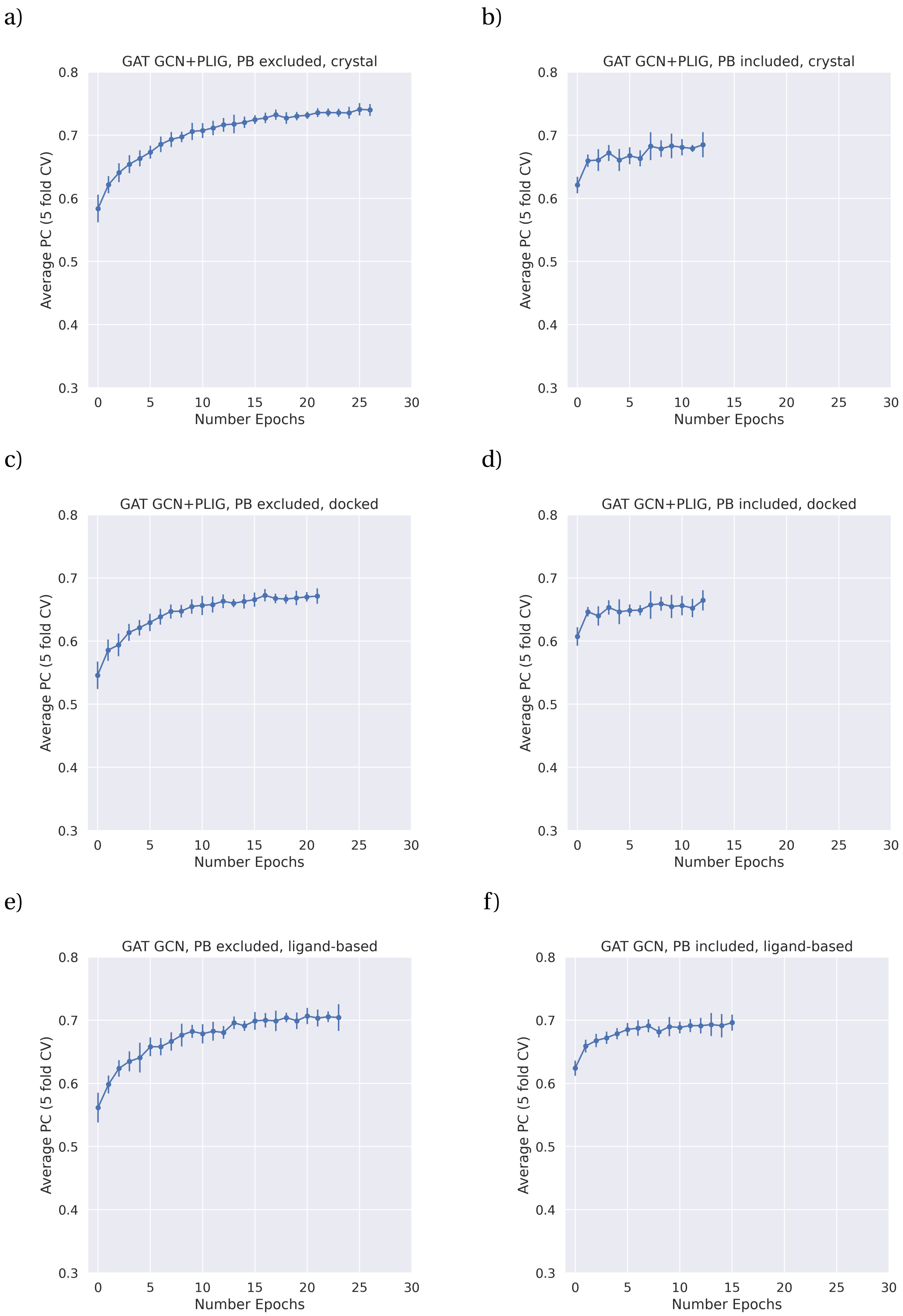

**Figure 3.4:** 5-fold cross validation performance of the GAT/GCN models reported as the average calculated pearson correlation coefficient and it's corresponding standard deviation (error bars) at each epoch. a) PB excluded, crystal structures; b) PB included, crystal structures; c) PB excluded, docked poses; d) PB included, docked poses, e) PB excluded, ligand-based, f) PB included, ligand-based.

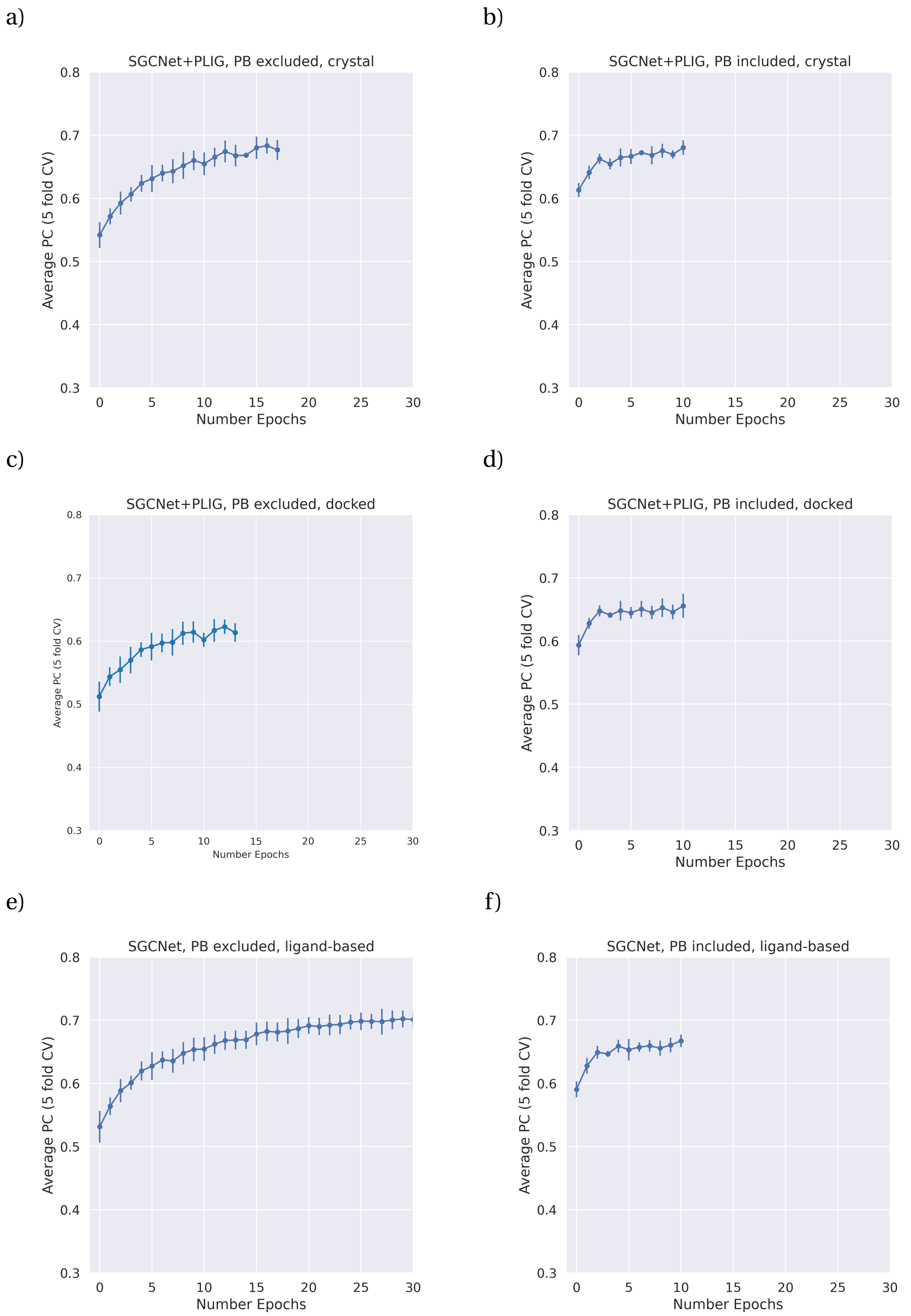

**Figure 3.5:** 5-fold cross validation performance of the SGCNet models reported as the average calculated pearson correlation coefficient and it's corresponding standard deviation (error bars) at each epoch. a) PB excluded, crystal structures; b) PB included, crystal structures; c) PB excluded, docked poses; d) PB included, docked poses, e) PB excluded, ligand-based, f) PB included, ligand-based.

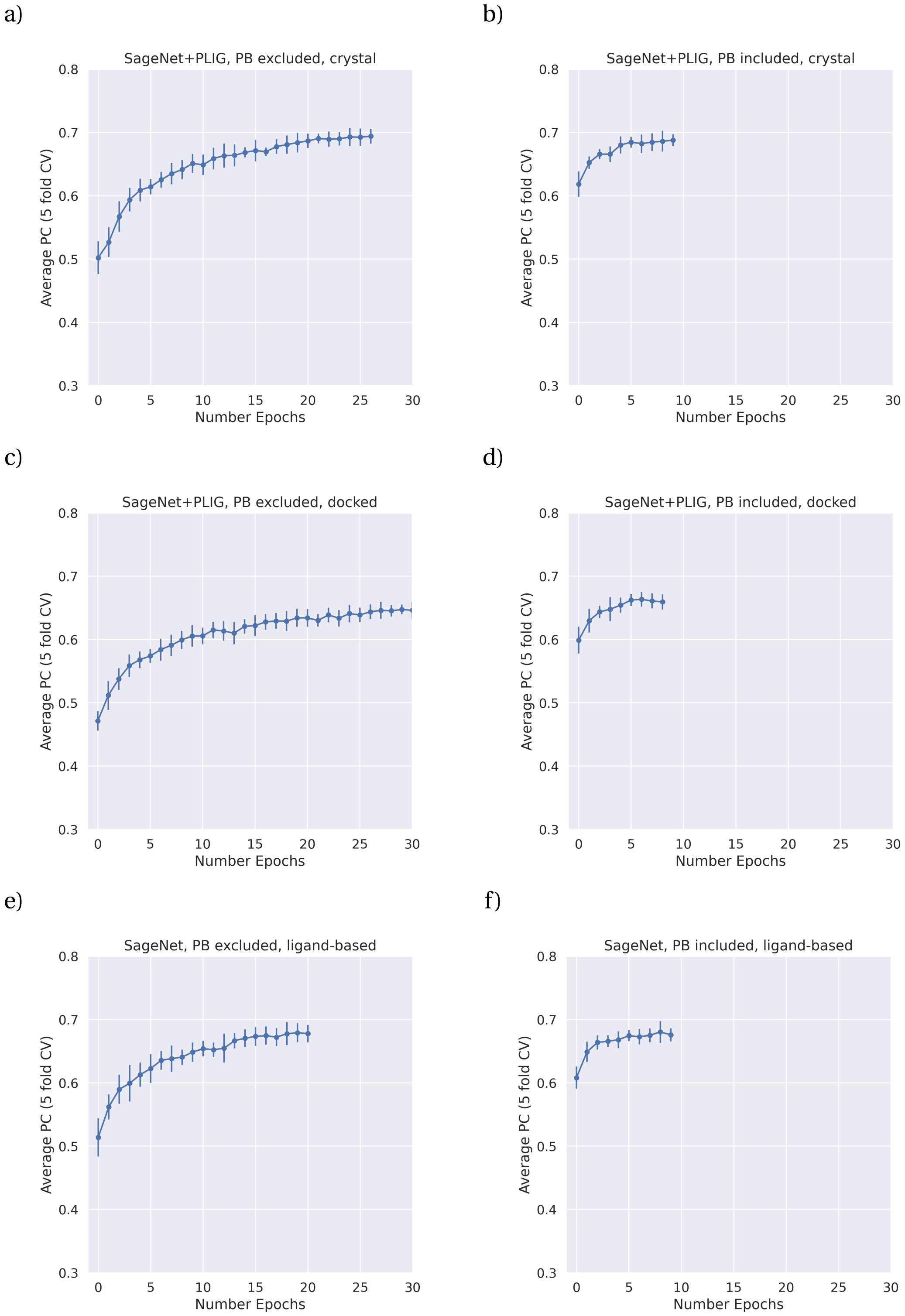

**Figure 3.6:** 5-fold cross validation performance of the Sage models reported as the average calculated pearson correlation coefficient and it's corresponding standard deviation (error bars) at each epoch. a) PB excluded, crystal structures; b) PB included, crystal structures; c) PB excluded, docked poses; d) PB included, docked poses, e) PB excluded, ligand-based, f) PB included, ligand-based.

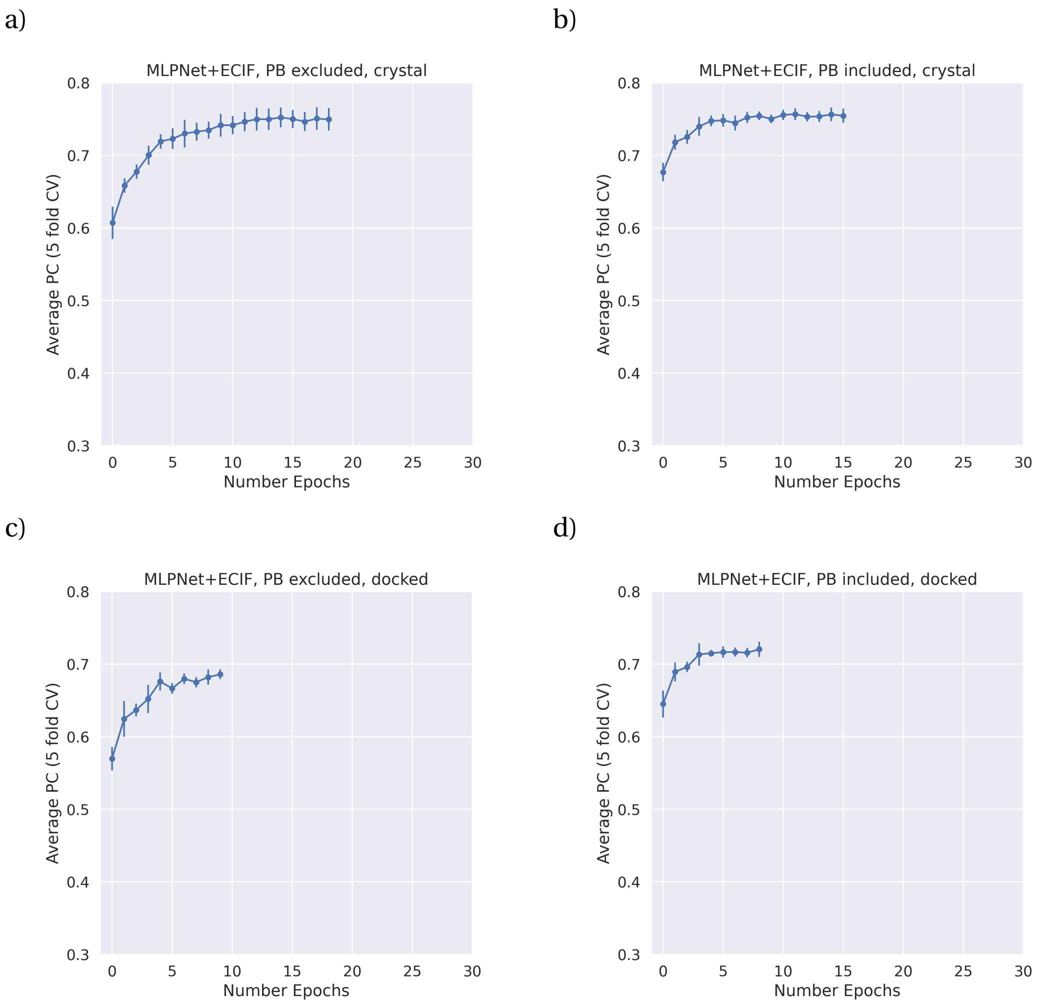

**Figure 3.7:** 5-fold cross validation performance of the MLPNet + ECIF models reported as the average calculated pearson correlation coefficient and it's corresponding standard deviation (error bars) at each epoch. a) PB excluded, crystal structures; b) PB included, crystal structures; c) PB excluded, docked poses; d) PB included, docked poses.

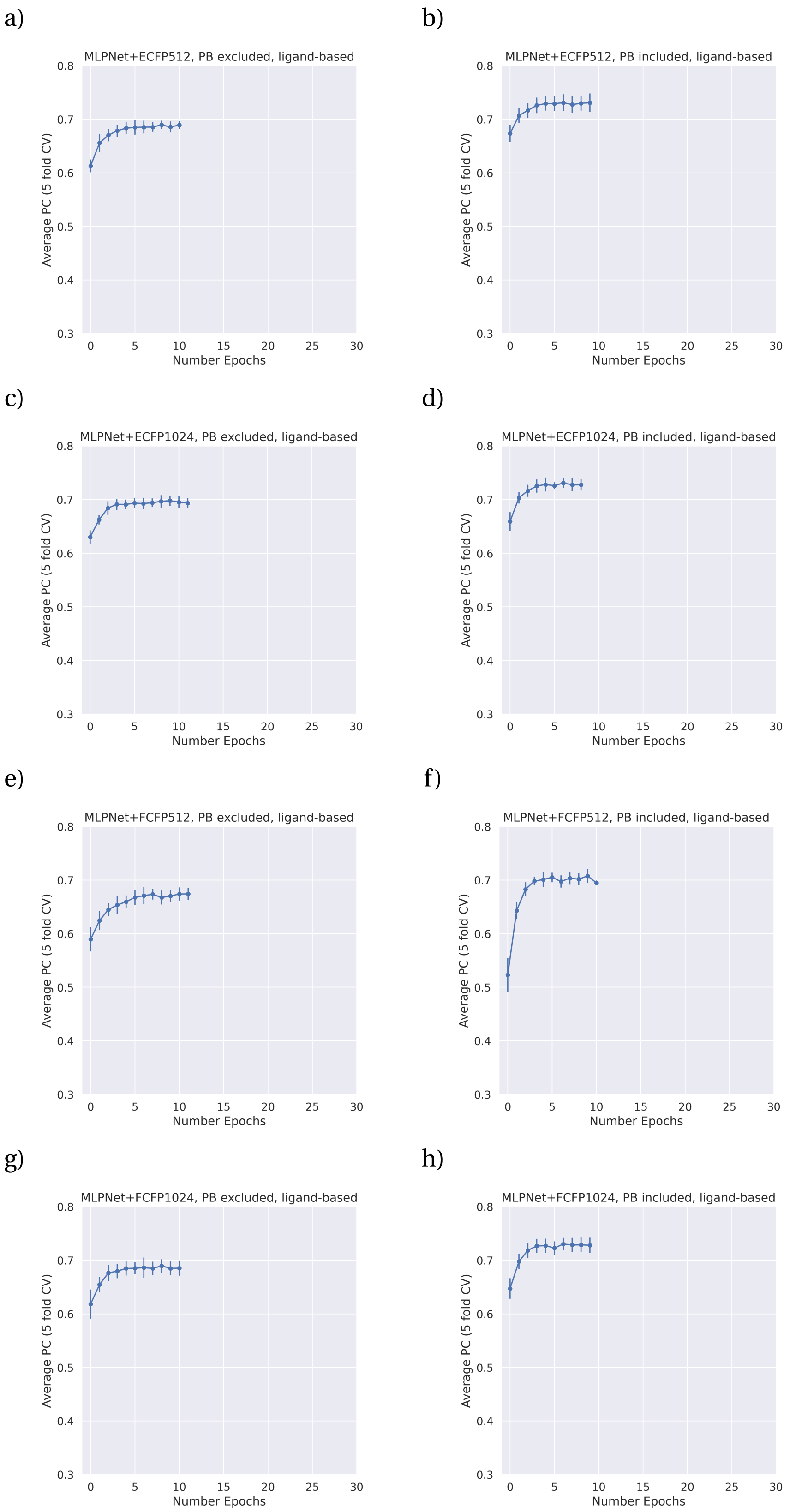

**Figure 3.8:** 5-fold cross validation performance of the MLPNet ligand-based models reported as the average calculated pearson correlation coefficient and it's corresponding standard deviation (error bars) at each epoch. a) ECFP512, PB excluded; b) ECFP512, PB included; c) ECFP1024, PB excluded; d) ECFP1024, PB included; e) FCFP512, PB excluded; f) FCFP512, PB included; g) FCFP1024, PB excluded; h) FCFP1024, PB included.

### 4. *Performance and stability of all models*

All models were trained on the PDBBind dataset (description see main text Section 2.1) and tested on the CASF-2016 benchmark set (crystal case is trained and tested on crystal structures, docked case is trained and tested on docked poses and the ligand case does not use 3D information). Since the models' predictions are somewhat stochastic, model performance and stability against the withheld test set (CASF-2016) for all trained models was evaluated using the average and standard deviation (SD) of the pearson correlation coefficient (PC) as well as the root-mean-square error (RMSE) over 10 runs. PC and RMSE and their corresponding standard deviations are shown in figure 4.1 and figure 4.2. The scatter plots of the average prediction versus the experimentally determined pK value for each protein-ligand complex between the 10 model runs for each model and feature combination are shown in Figure 4.3-4.11. The best performing PLIG model was the GATNet PLIG without protein sequence embedding (PC = 0.80) and the best performing model overall was the MLPNet ECIF model with protein sequence embedding (PC = 0.82). However, since the standard deviation of the Pearson correlation coefficient is as large as the difference between the GATNet PLIG and MLPNet ECIF model (Figure 4.1, standard deviation of 0.018 versus 0.009 for GATNet PLIG and MLPNet ECIF, respectively), the difference is not significant and both models should be considered to be of similar performance. Overall, model stability as measured by PC standard deviation varied between 0.007 (GCNNet PLIG, no sequence trained and tested on crystal poses) and 0.023 (SGCNet ligand-based, no sequence)

In addition, the case where models were trained on crystal poses and tested on docked poses was tested as well to investigate if performance improves. The results are shown in Figure 4.12. There is no significant difference when training on crystal poses and testing on docked poses as opposed to the standard case (trained and tested on docked poses).

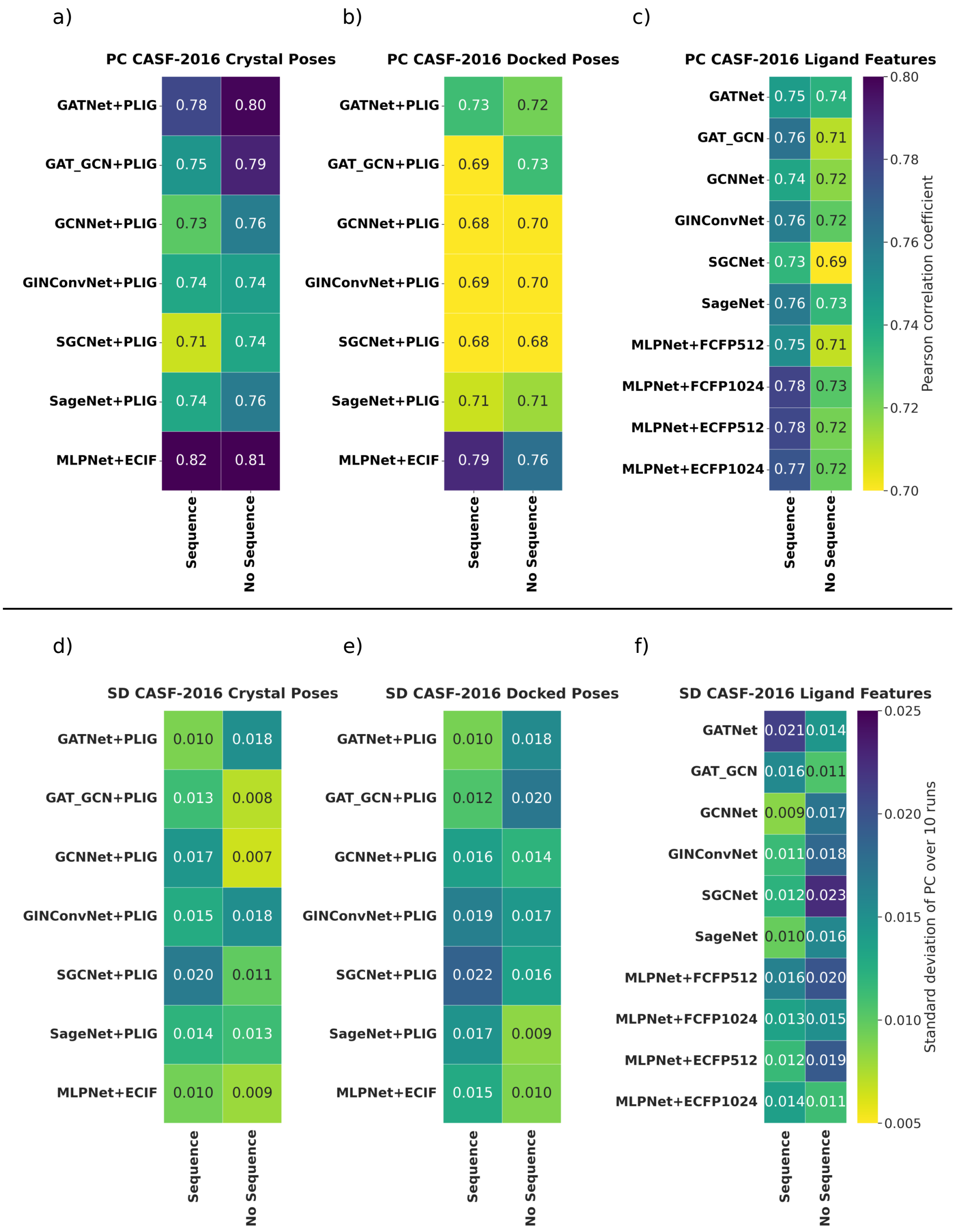

**Figure 4.1:** The reported Pearson correlation coefficient (PC) is calculated as the average over 10 train and test model iterations and shown in a-c with the corresponding standard deviation shown in d-f. *Sequence* and *No Sequence* denote the presence or absence of the protein sequence encoding branch in the model architecture, respectively. (a and d) PC and SD of the structure-based GNN and MLPNet models when trained and tested on crystal structures. (b and e) PC and SD of the structure-based GNN and MLPNet models when trained and tested on docked poses. (c and f) PC and SD of all ligand-based GNN and MLPNet models. ECFP and FCFP fingerprint radius was 2 in all cases. Best performing PLIG model was the GATNet PLIG without protein sequence embedding (PC = 0.80) and the best performing model overall was the MLPNet ECIF model with protein sequence embedding (PC = 0.82). Model stability as measured by PC standard deviation varied between 0.007 (GCNNet PLIG no sequence trained/tested on crystal poses) and 0.023 (SGCNet ligand-based no sequence).

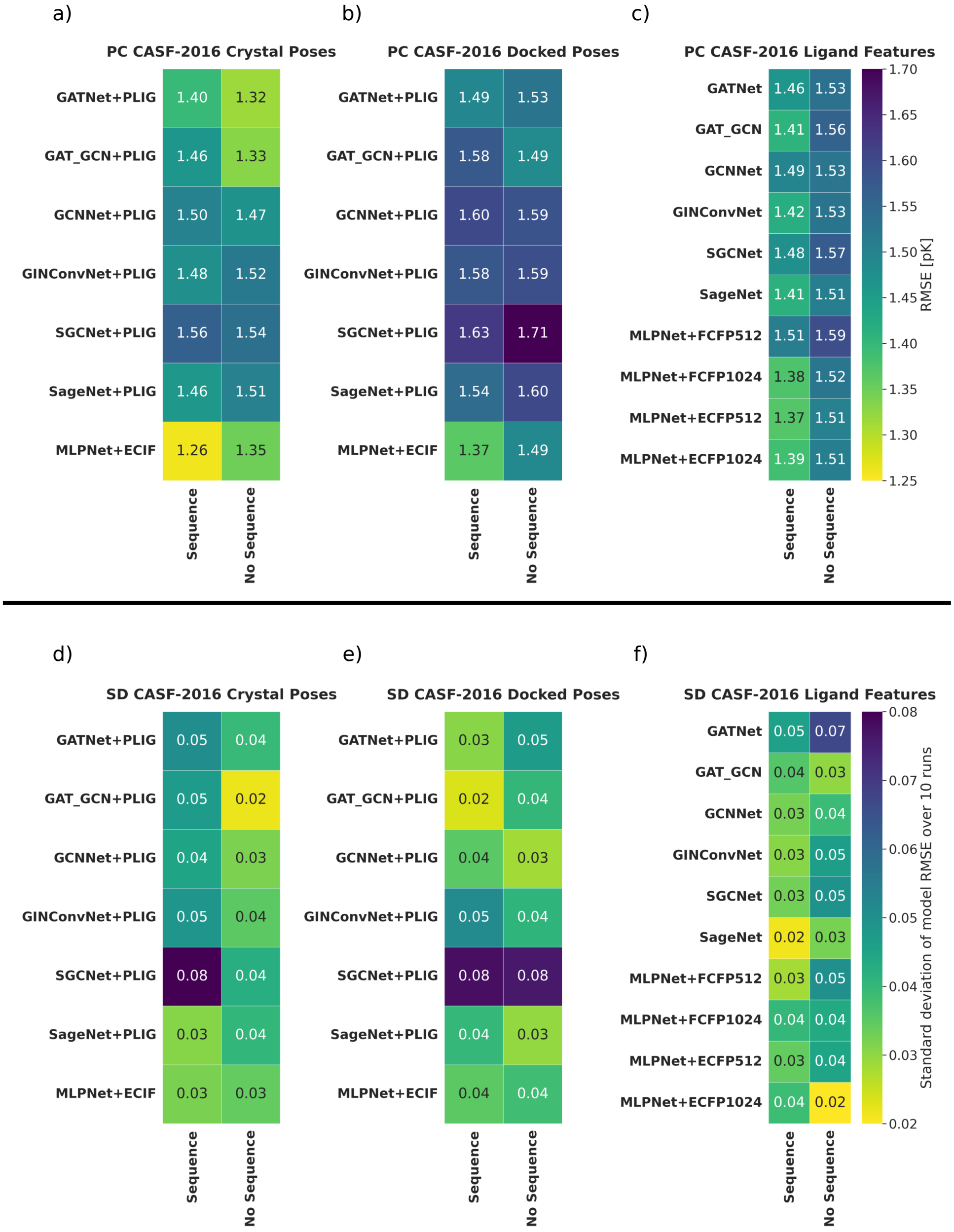

**Figure 4.2:** The reported root-mean-square error (RMSE) is calculated as the average over 10 train and test model iterations and shown in a-c with the corresponding standard deviation shown in d-f. *Sequence* and *No Sequence* denote the presence or absence of the protein sequence encoding branch in the model architecture, respectively. (a and d) RMSE and SD of the structure-based GNN and MLPNet models when trained and tested on crystal structures. (b and e) RMSE and SD of the structure-based GNN and MLPNet models when trained and tested on docked poses. (c and f) RMSE and SD of all ligand-based GNN and MLPNet models. ECFP and FCFP fingerprint radius was 2 in all cases. Best performing PLIG model was the GATNet PLIG without protein sequence embedding (RMSE = 1.32) and the best performing model overall was the MLPNet ECIF model with protein sequence embedding (RMSE = 1.26). Model stability as measured by RMSE standard deviation varied between 0.02 (several models) and 0.08 (several SGCNet based models).

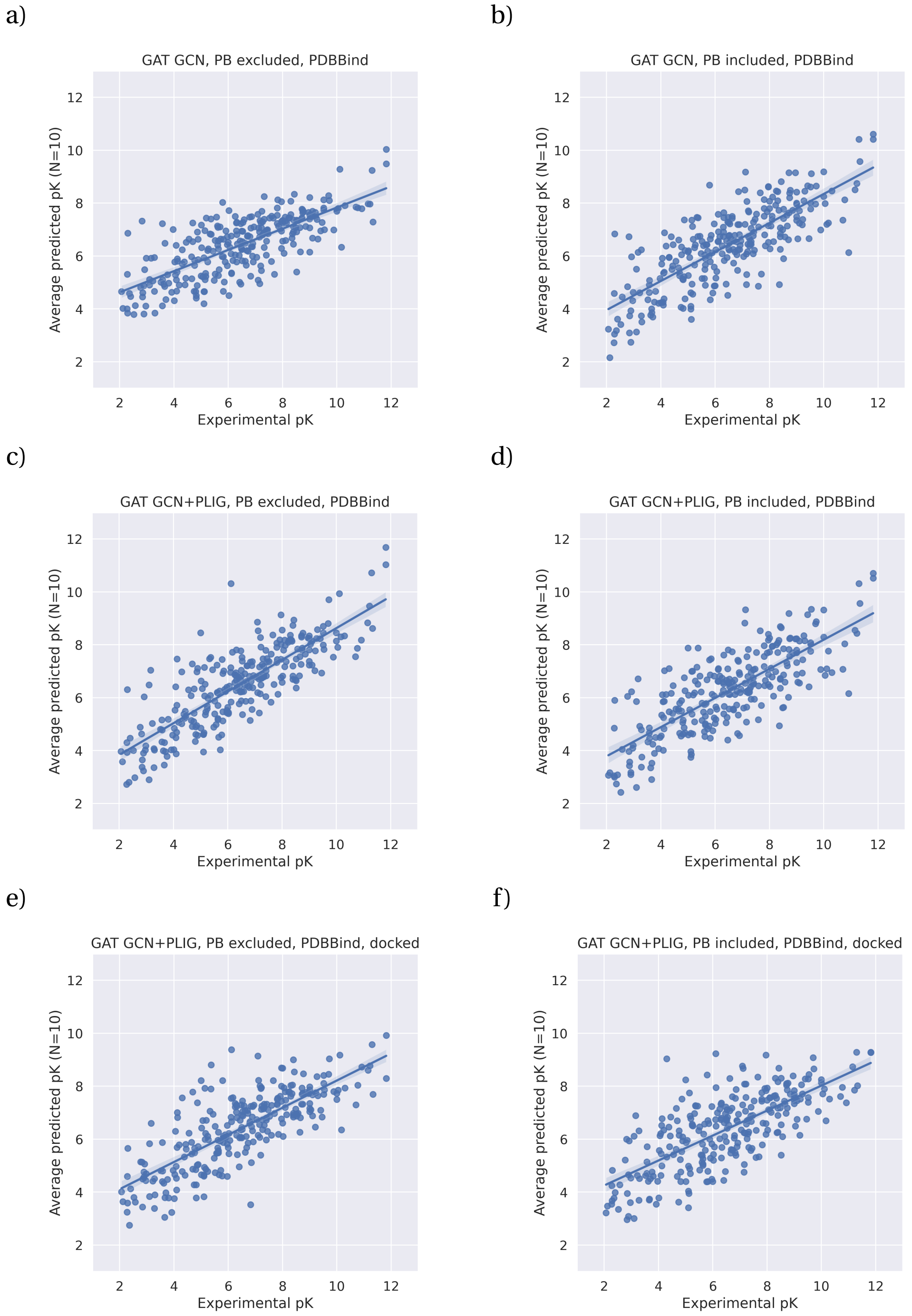

**Figure 4.3:** Scatter plots of the average prediction versus the experimentally determined pK value for each protein-ligand complex between the 10 model runs for all GAT+GCN models. a) ligand-based graph, no protein sequence embedding; b) ligand-based graph, including protein embedding; c) PLIG, no protein sequence embedding, trained and tested on crystal structures; d) PLIG, including sequence embedding, trained and tested on crystal structures; e) PLIG, no protein sequence embedding, trained and tested on docked structures; f) PLIG, including protein sequence embedding, trained and tested on docked structures.

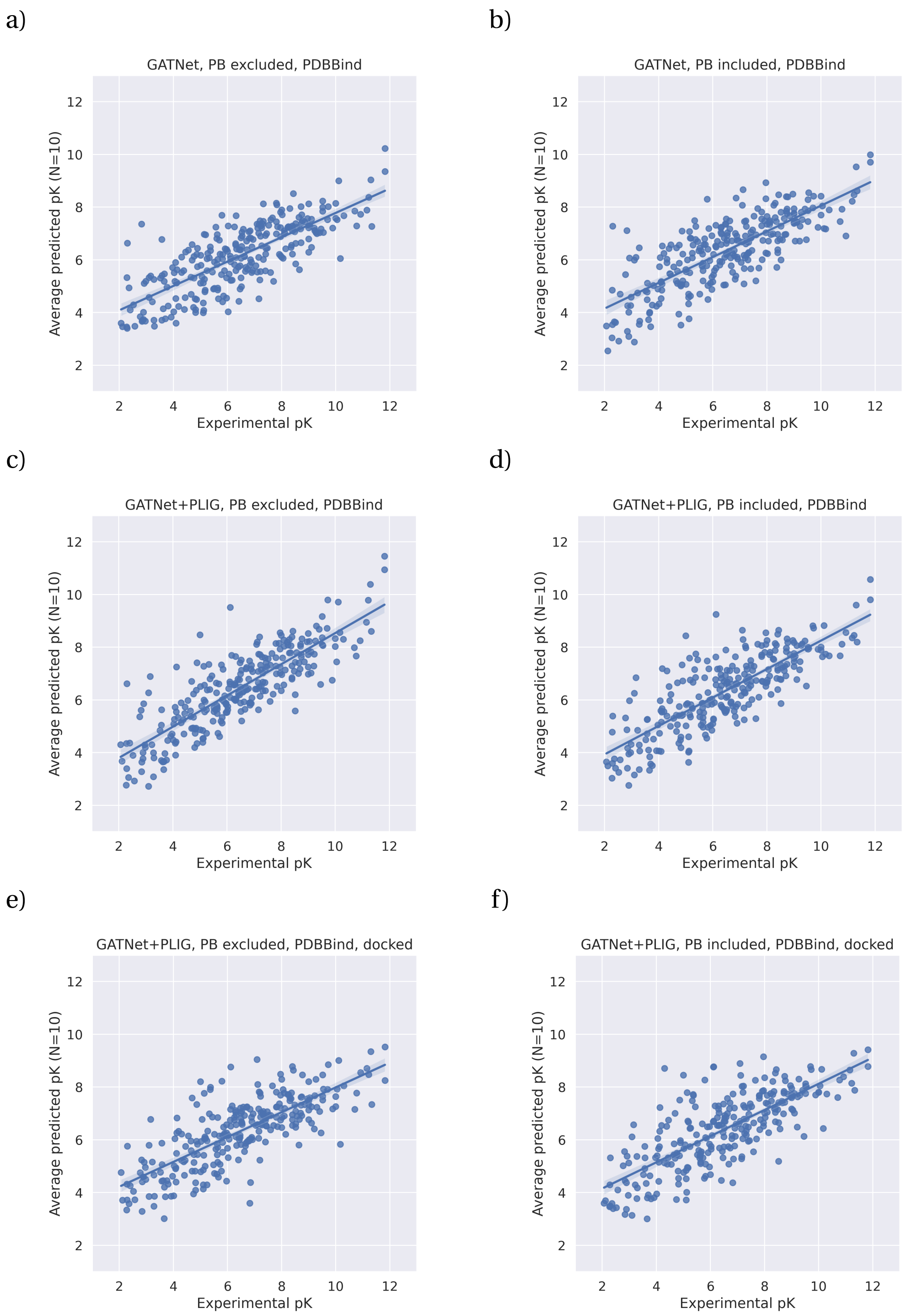

**Figure 4.4:** Scatter plots of the average prediction versus the experimentally determined pK value for each protein-ligand complex between the 10 model runs for all GATNet models. a) ligand-based graph, no protein sequence embedding; b) ligand-based graph, including protein embedding; c) PLIG, no protein sequence embedding, trained and tested on crystal structures; d) PLIG, including sequence embedding, trained and tested on crystal structures; e) PLIG, no protein sequence embedding, trained and tested on docked structures; f) PLIG, including protein sequence embedding, trained and tested on docked structures.

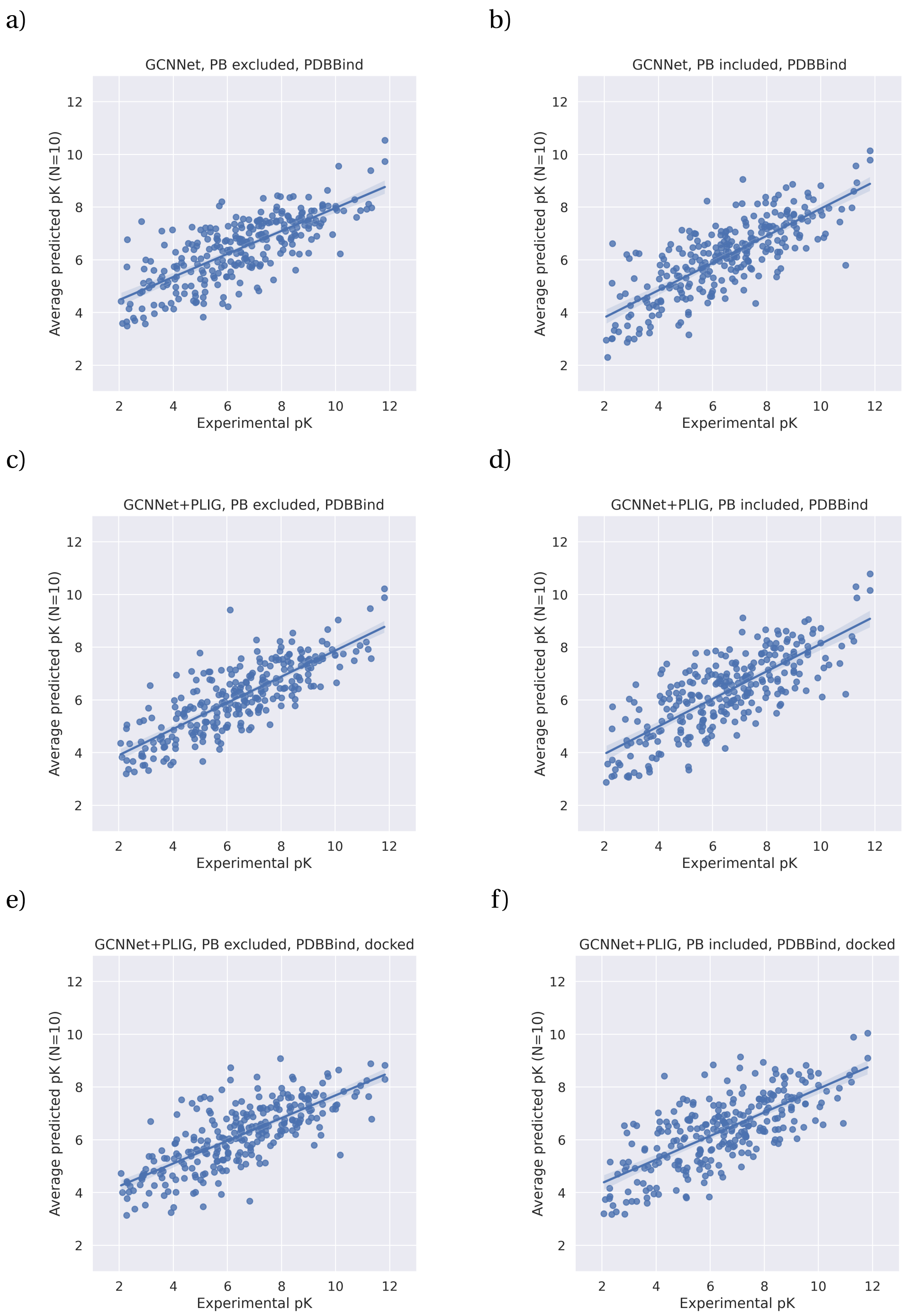

**Figure 4.5:** Scatter plots of the average prediction versus the experimentally determined pK value for each protein-ligand complex between the 10 model runs for all GCNNet models. a) ligand-based graph, no protein sequence embedding; b) ligand-based graph, including protein embedding; c) PLIG, no protein sequence embedding, trained and tested on crystal structures; d) PLIG, including sequence embedding, trained and tested on crystal structures; e) PLIG, no protein sequence embedding, trained and tested on docked structures; f) PLIG, including protein sequence embedding, trained and tested on docked structures.

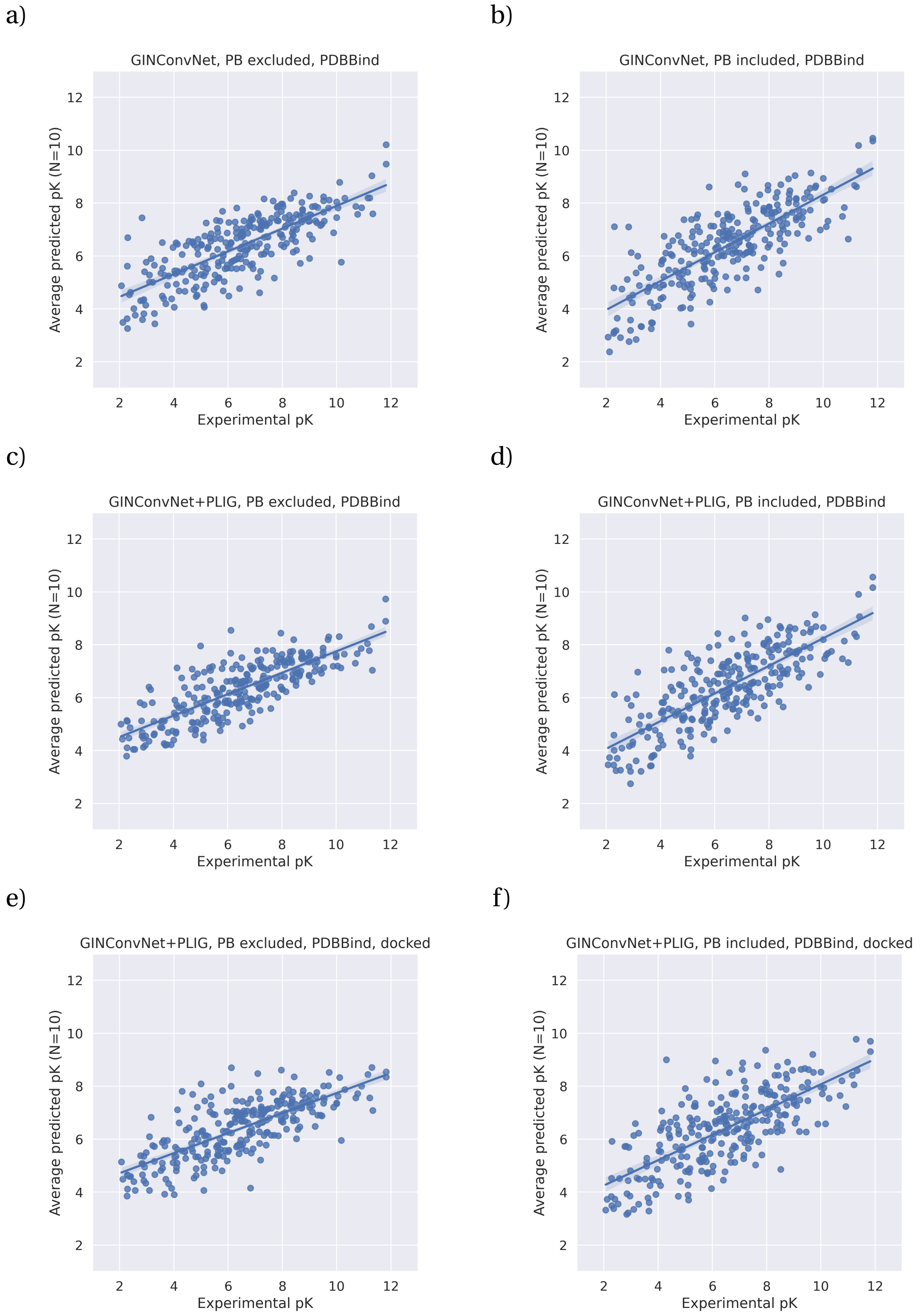

**Figure 4.6:** Scatter plots of the average prediction versus the experimentally determined pK value for each protein-ligand complex between the 10 model runs for all GinConvNet models. a) ligand-based graph, no protein sequence embedding; b) ligand-based graph, including protein embedding; c) PLIG, no protein sequence embedding, trained and tested on crystal structures; d) PLIG, including sequence embedding, trained and tested on crystal structures; e) PLIG, no protein sequence embedding, trained and tested on docked structures; f) PLIG, including protein sequence embedding, trained and tested on docked structures.

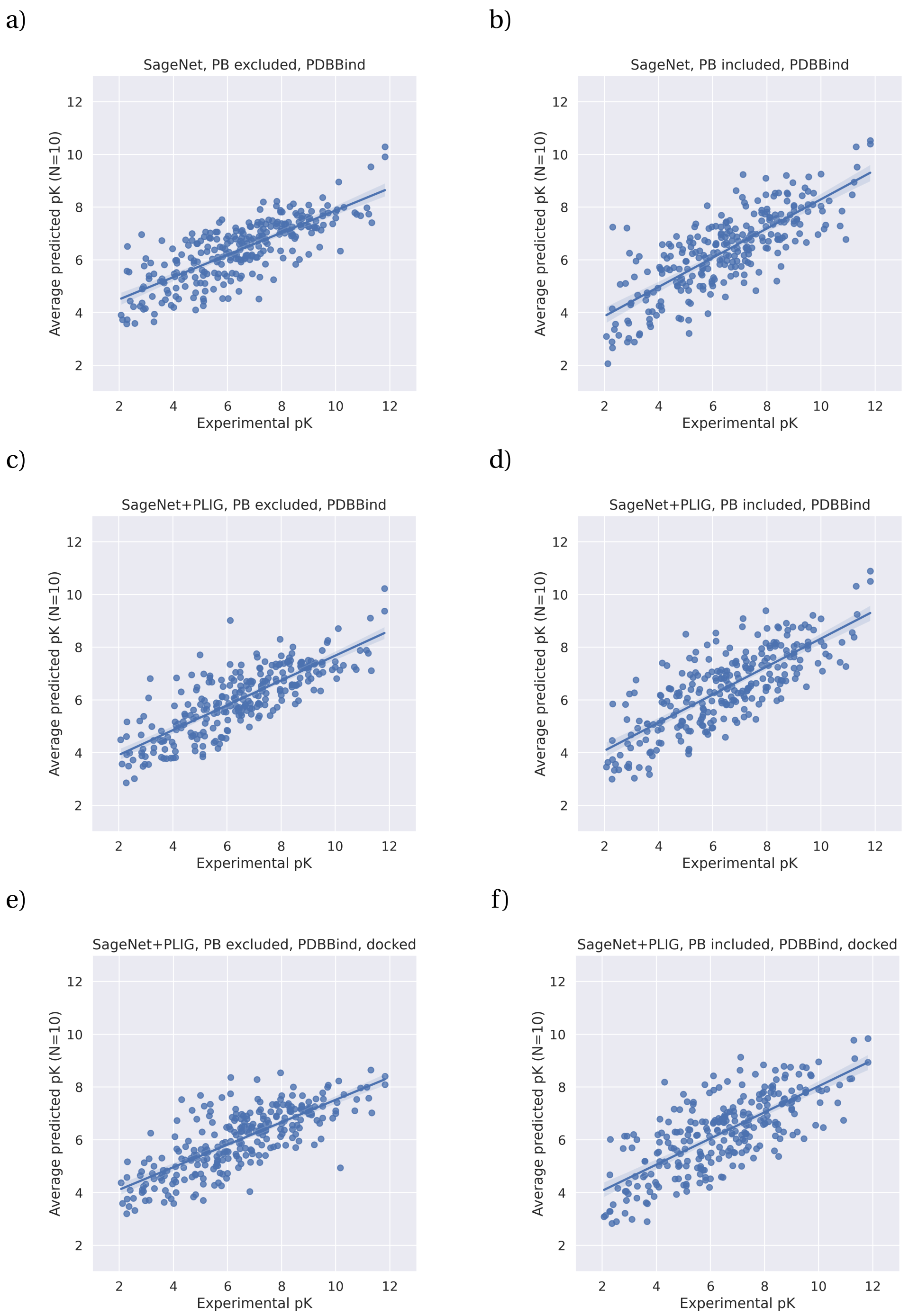

**Figure 4.7:** Scatter plots of the average prediction versus the experimentally determined pK value for each protein-ligand complex between the 10 model runs for all SageNet models. a) ligand-based graph, no protein sequence embedding; b) ligand-based graph, including protein embedding; c) PLIG, no protein sequence embedding, trained and tested on crystal structures; d) PLIG, including sequence embedding, trained and tested on crystal structures; e) PLIG, no protein sequence embedding, trained and tested on docked structures; f) PLIG, including protein sequence embedding, trained and tested on docked structures.

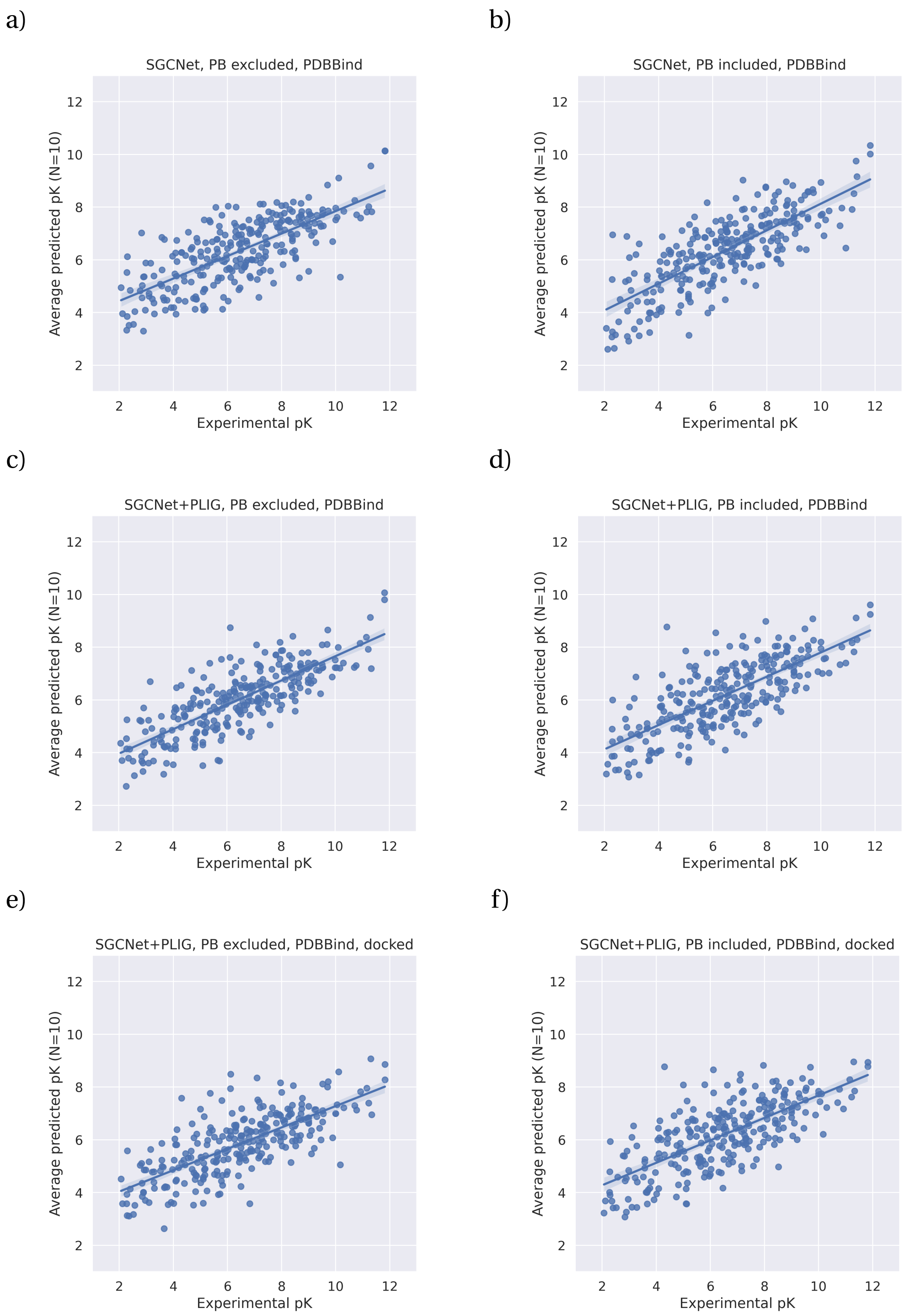

**Figure 4.8:** Scatter plots of the average prediction versus the experimentally determined pK value for each protein-ligand complex between the 10 model runs for all SGCNet models. a) ligand-based graph, no protein sequence embedding; b) ligand-based graph, including protein embedding; c) PLIG, no protein sequence embedding, trained and tested on crystal structures; d) PLIG, including sequence embedding, trained and tested on crystal structures; e) PLIG, no protein sequence embedding, trained and tested on docked structures; f) PLIG, including protein sequence embedding, trained and tested on docked structures.

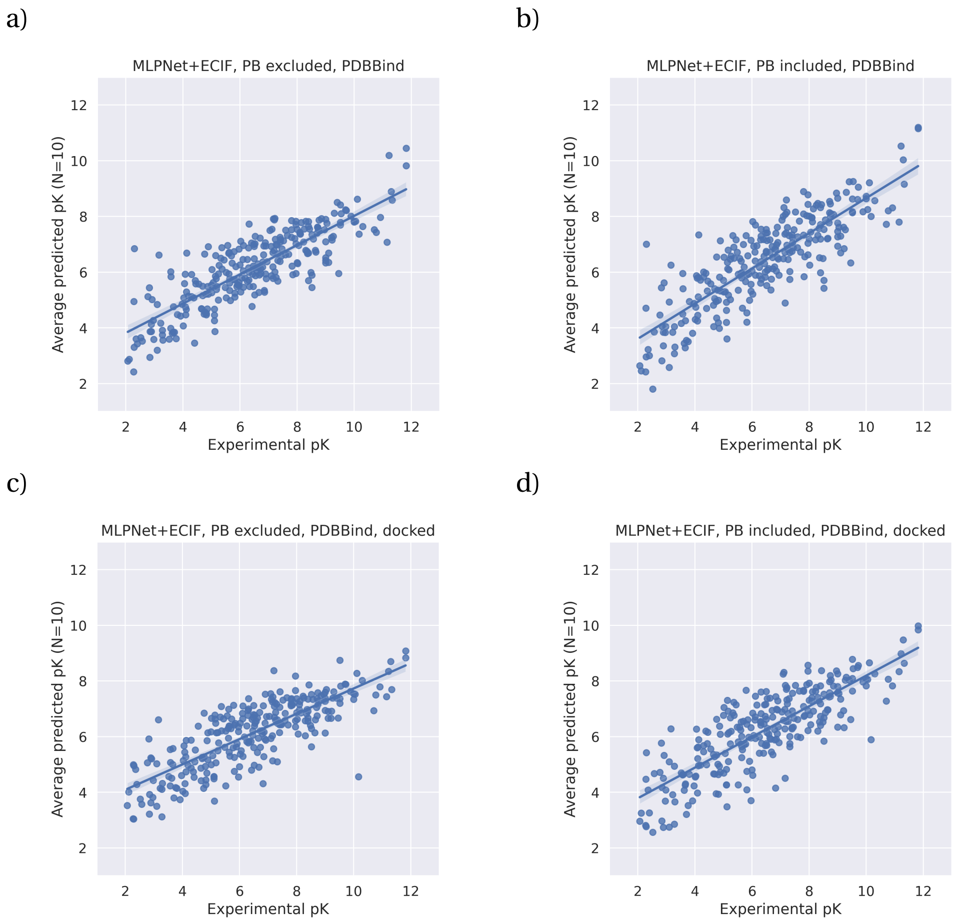

**Figure 4.9:** Scatter plots of the average prediction versus the experimentally determined pK value for each protein-ligand complex between the 10 model runs for all MLPNet ECIF models. a) MLPNet ECIF, no protein sequence embedding, trained and tested on crystal structures; b) MLPNet ECIF, including sequence embedding, trained and tested on crystal structures; c) MLPNet ECIF, no protein sequence embedding, trained and tested on docked structures; d) MLPNet ECIF, including protein sequence embedding, trained and tested on docked structures.

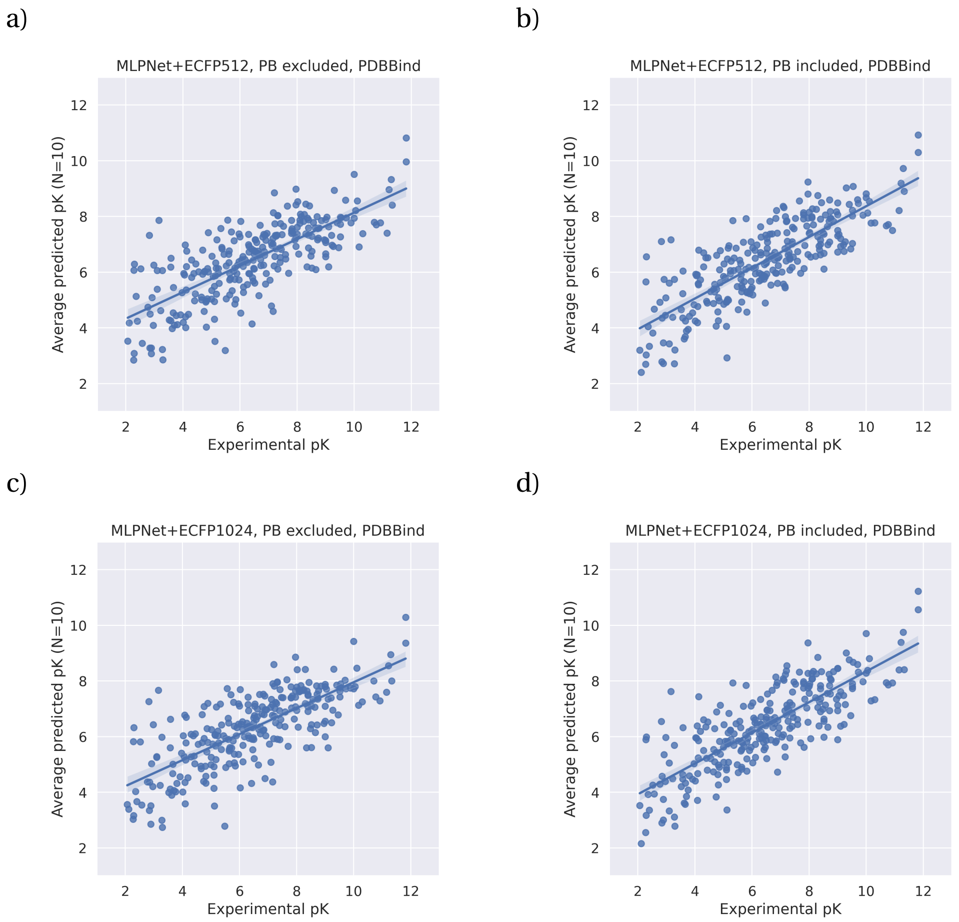

**Figure 4.10:** Scatter plots of the average prediction versus the experimentally determined pK value for each protein-ligand complex between the 10 model runs for all MLPNet ECFP models. a) MLPNet ECFP512, no protein sequence embedding, trained and tested on ligand information; b) MLPNet ECFP512, including sequence embedding, trained and tested on ligand information; c) MLPNet ECFP1024, no protein sequence embedding, trained and tested on ligand information; d) MLPNet ECFP1024, including protein sequence embedding, trained and tested on ligand information

a)

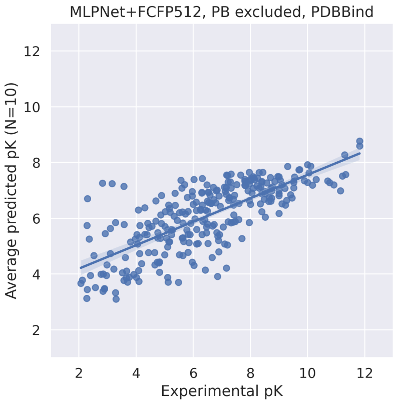

b)

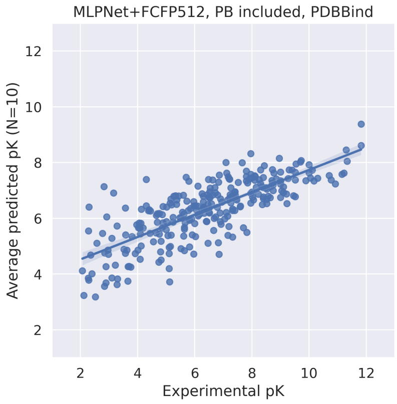

c)

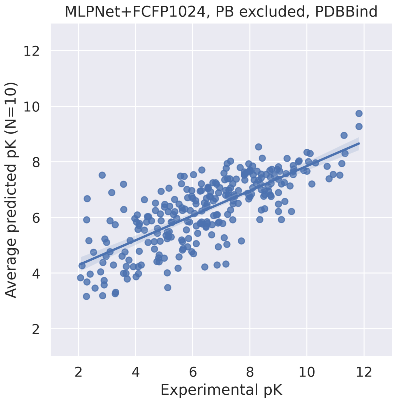

d)

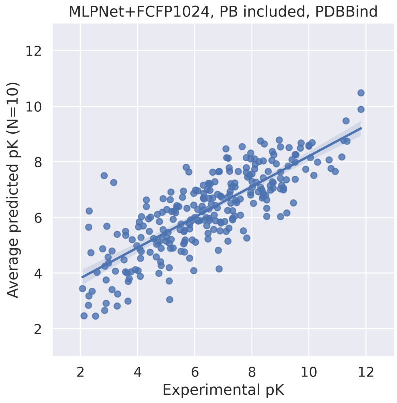

**Figure 4.11:** Scatter plots of the average prediction versus the experimentally determined pK value for each protein-ligand complex between the 10 model runs for all MLPNet FCFP models. a) MLPNet FCFP512, no protein sequence embedding, trained and tested on ligand information; b) MLPNet FCFP512, including sequence embedding, trained and tested on ligand information; c) MLPNet FCFP1024, no protein sequence embedding, trained and tested on ligand information; d) MLPNet FCFP1024, including protein sequence embedding, trained and tested on ligand information

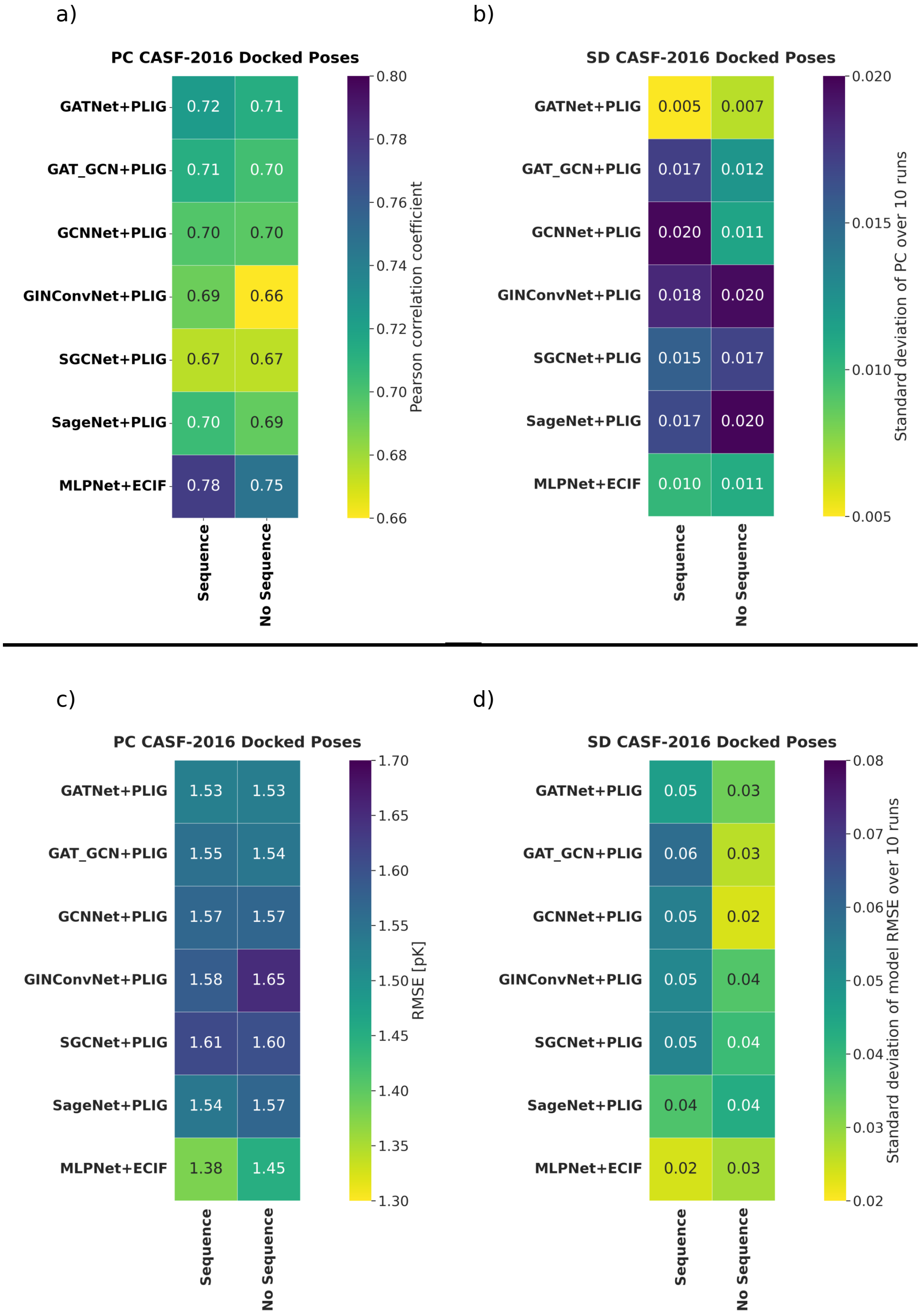

**Figure 4.12:** The reported Pearson correlation coefficient (PC) and Root-mean-square error (RMSE) is calculated as the average over 10 train and test model iterations with the corresponding standard deviation. *Sequence* and *No Sequence* denote the presence or absence of the protein sequence encoding branch in the model architecture, respectively. (a and b) PC and SD of the structure-based GNN and MLPNet models when trained on crystal poses and tested on docked poses. (c and d) RMSE and SD of the structure-based GNN and MLPNet models when trained on crystal poses and tested on docked poses. Best performing model overall was the MLPNet ECIF model with protein sequence embedding (PC = 0.78, RMSE = 1.38). There is no significant performance difference to models trained and tested on docked poses as shown in Figure 4.1 amd 4.2

### 5. *Proximity Analysis*

#### 5.1 Cross validation

5-fold cross validation was performed on the combined PDBBind general 2020 and PDBBind refined 2016 set (dataset details in the main text Section 2.1). The validation and test set (CASF-2016) used in the main study was removed from the cross validation set, to leave the training set of 14254 compounds. Cross validation was done separately for the docked and crystal-based datasets to ensure model stability for both, crystal derived structures and docked poses. This set was split into 5 random folds, using 20 % of the dataset as validation in each fold. Models were run across all 5 folds until model performance converged. The performance of all models during cross validation is recorded for every epoch. The number of epochs to train each model for training and test on the CASF-2016 benchmark was determined as the point where no significant performance increase (difference of less than 0.01 Pearson correlation coefficient between epoch $n$ and $n-1$). The GATNet model was chosen without the protein sequence embedding branch as the architecture to try out different PLIG proximity thresholds since it was the best performing model in the main study. The optimal number of epochs as determined by cross validation is shown in table 5.1.

| **Proximity threshold** | **Crystal** | **Docked** |
|---|---|---|
| PLIG 4 Å | 12 | 11 |
| PLIG 5 Å | 14 | 9 |
| PLIG 6 Å | 16 | 11 |
| PLIG 7 Å | 14 | 15 |
| PLIG 8 Å | 15 | 13 |

**Table 5.1:** Optimal number of epochs for different PLIG thresholds when trained and tested on docked and crystal poses using the GATNet PLIG architecture without the protein sequence embedding.

The following section shows the performance of each of the 4,5,7 and 8 Å models over every epoch used for the determination of the optimal number of epochs. The 6 Å PLIG model can be found in the general cross validations Section 3 above.

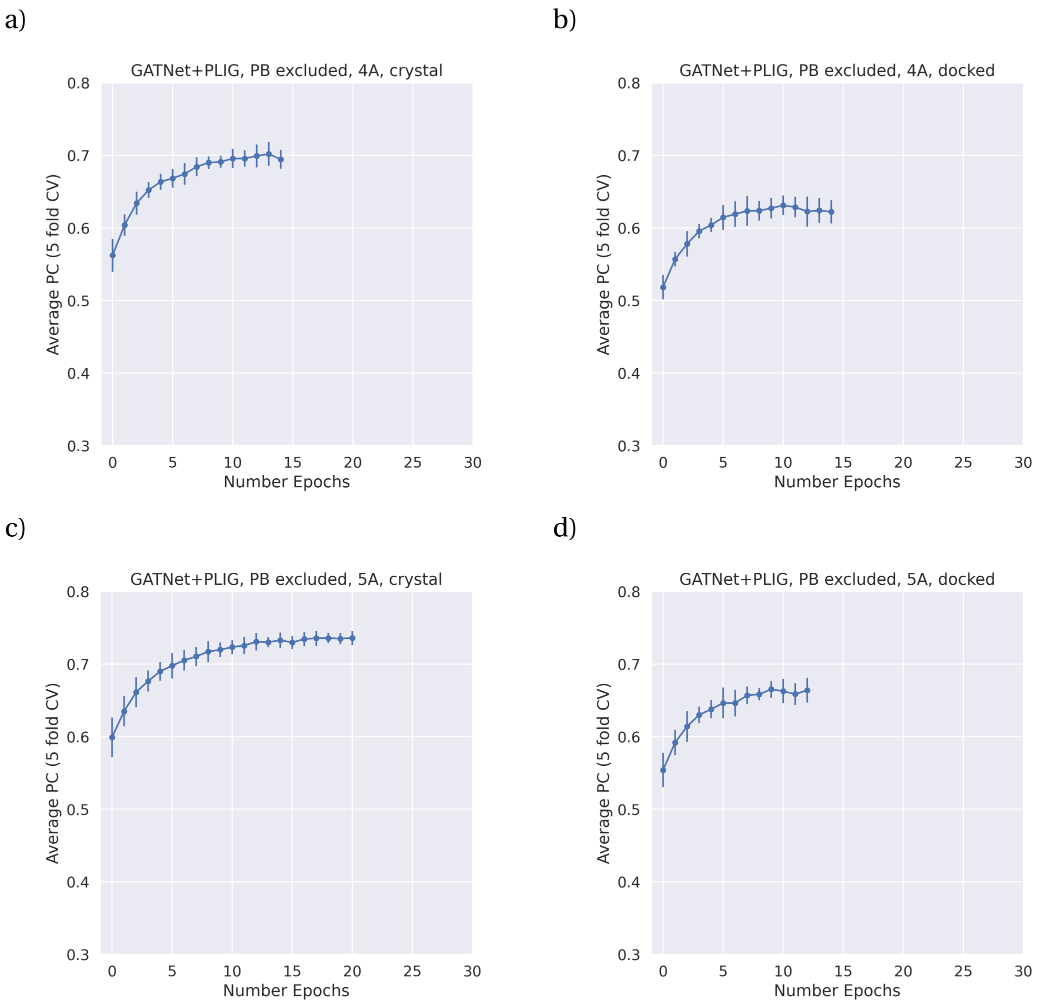

**Figure 5.1:** 5-fold cross validation performance of the GATNet PLIG models (no sequence) with 4 and 5 Å proximity thresholds reported as the average calculated pearson correlation coefficient and it's corresponding standard deviation (error bars) at each epoch. a) 4 Å threshold, trained and tested on crystal poses; b) 4 Å threshold, trained and tested on docked poses; c) 5 Å threshold, trained and tested on crystal poses ; d) 5 Å threshold, trained and tested on docked poses.

a)

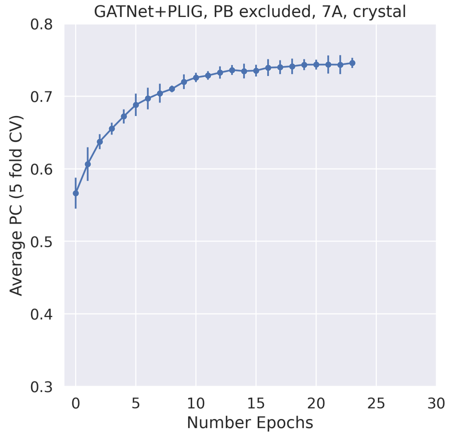

b)

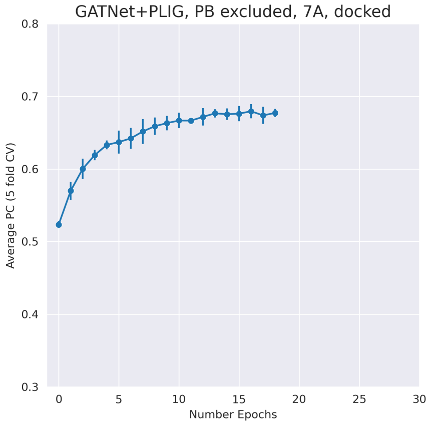

c)

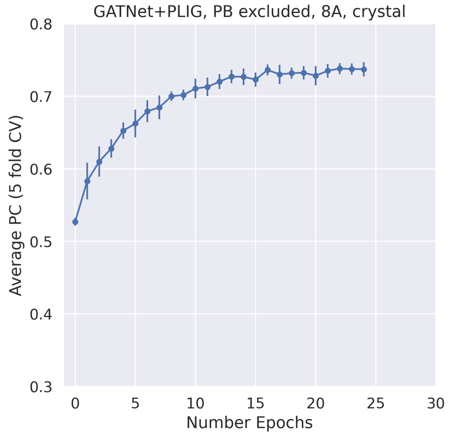

d)

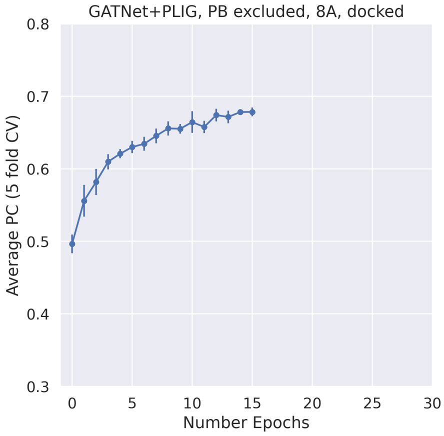

**Figure 5.2:** 5-fold cross validation performance of the GATNet PLIG models (no sequence) with 7 and 8 Å proximity thresholds reported as the average calculated pearson correlation coefficient and it's corresponding standard deviation (error bars) at each epoch. a) 7 Å threshold, trained and tested on crystal poses; b) 7 Å threshold, trained and tested on docked poses; c) 8 Å threshold, trained and tested on crystal poses ; d) 8 Å threshold, trained and tested on docked poses.

#### 5.2 Performance and stability over 10 stochastic runs

All models were trained on the PDBBind dataset (description see main text Section 2.1) and tested on the CASF-2016 benchmark set (crystal case is trained and tested on crystal structures, docked case is trained and tested on docked poses and the ligand case does not use 3D information). Since the models' predictions are somewhat stochastic, model performance and stability against the withheld test set (CASF-2016) for all trained models was evaluated using the average and standard deviation (SD) of the pearson correlation coefficient (PC) as well as the root-mean-square error (RMSE) over 10 independent runs. PC and RMSE and their corresponding standard deviations are shown in figure 5.3 and figure 5.4. Overall, the 5 and 6 Å representations reach similar performance metrics (PC =

0.80 / 0.80, RMSE = 1.33 / 1.32 for 5 / 6 Å respectively) however the 6 Å model is slightly more stable with a lower standard deviation for the RMSE (SD RMSE = 0.06 / 0.04 for 5 / 6 Å respectively). The difference in PC and standard deviation of the PC is insignificant between the two models. In addition, performance of all models against docked poses is extremely similar and overall lower than the crystal poses. This observation is in line with our expectations since a majority of docked poses were found to have a RMSD between the crystal and docked pose of larger than 2 Å (Main text Section 3.1). A threshold change of the same magnitude should therefore not alter results drastically, as most poses have an equally large inaccuracy in their pose.

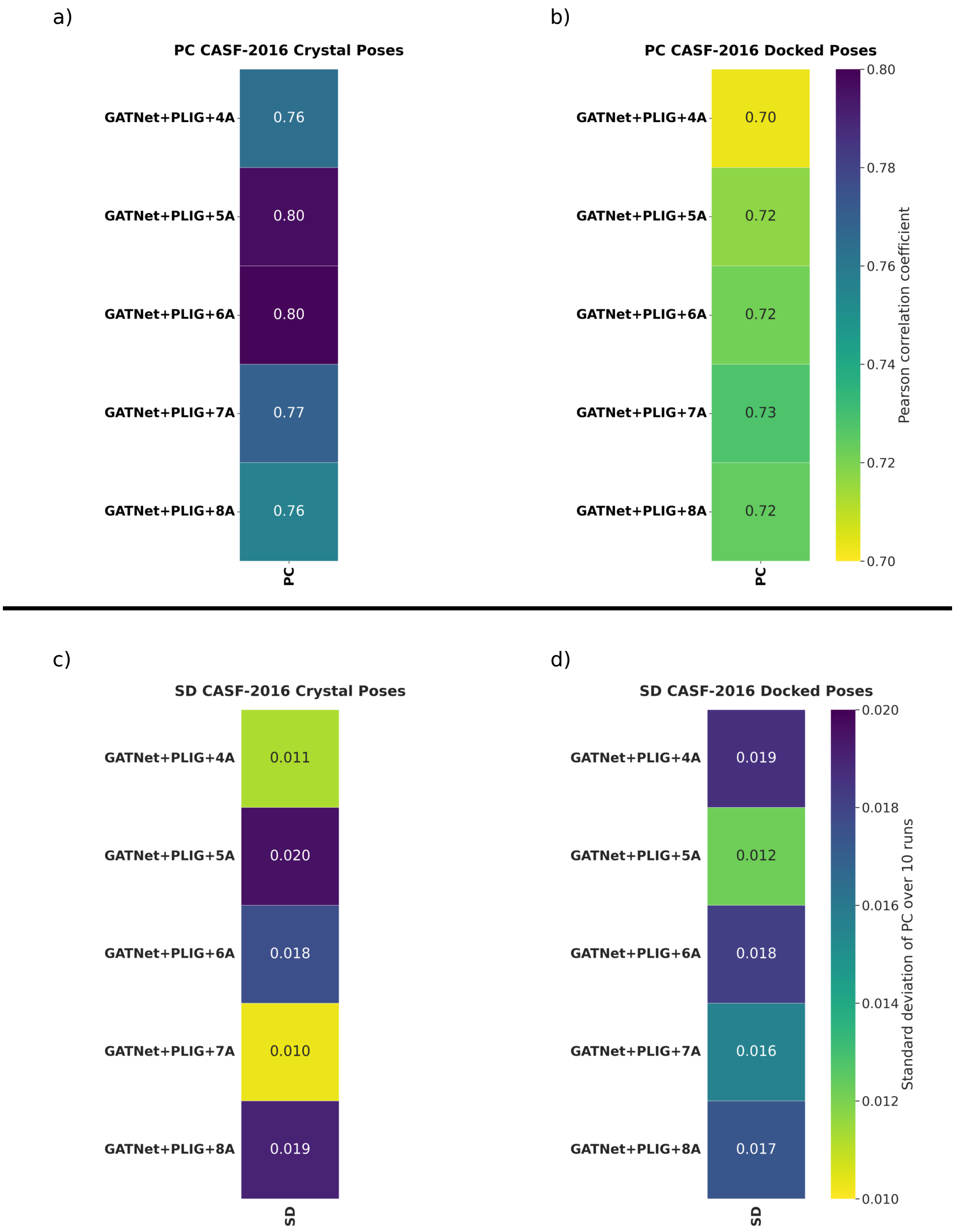

**Figure 5.3:** The reported Pearson correlation coefficient (PC) is calculated as the average over 10 train and test model iterations and shown in a-b with the corresponding standard deviation shown in c-d. All GATNet models were used without the second protein sequence embedding branch implementation. (a and c) PC and SD of the GATNet PLIG models with different proximity thresholds when trained and tested on crystal structures. (b and d) PC and SD of the GATNet PLIG models with different proximity thresholds when trained and tested on docked poses. Best performing threshold was the GATNet PLIG 5 and 6 Å models (PC = 0.80). Model stability as measured by PC standard deviation varied between 0.01 (GATNet PLIG 7 Å trained/tested on crystal poses) and 0.02 (GATNet PLIG 5 Å, trained/tested on crystal poses).

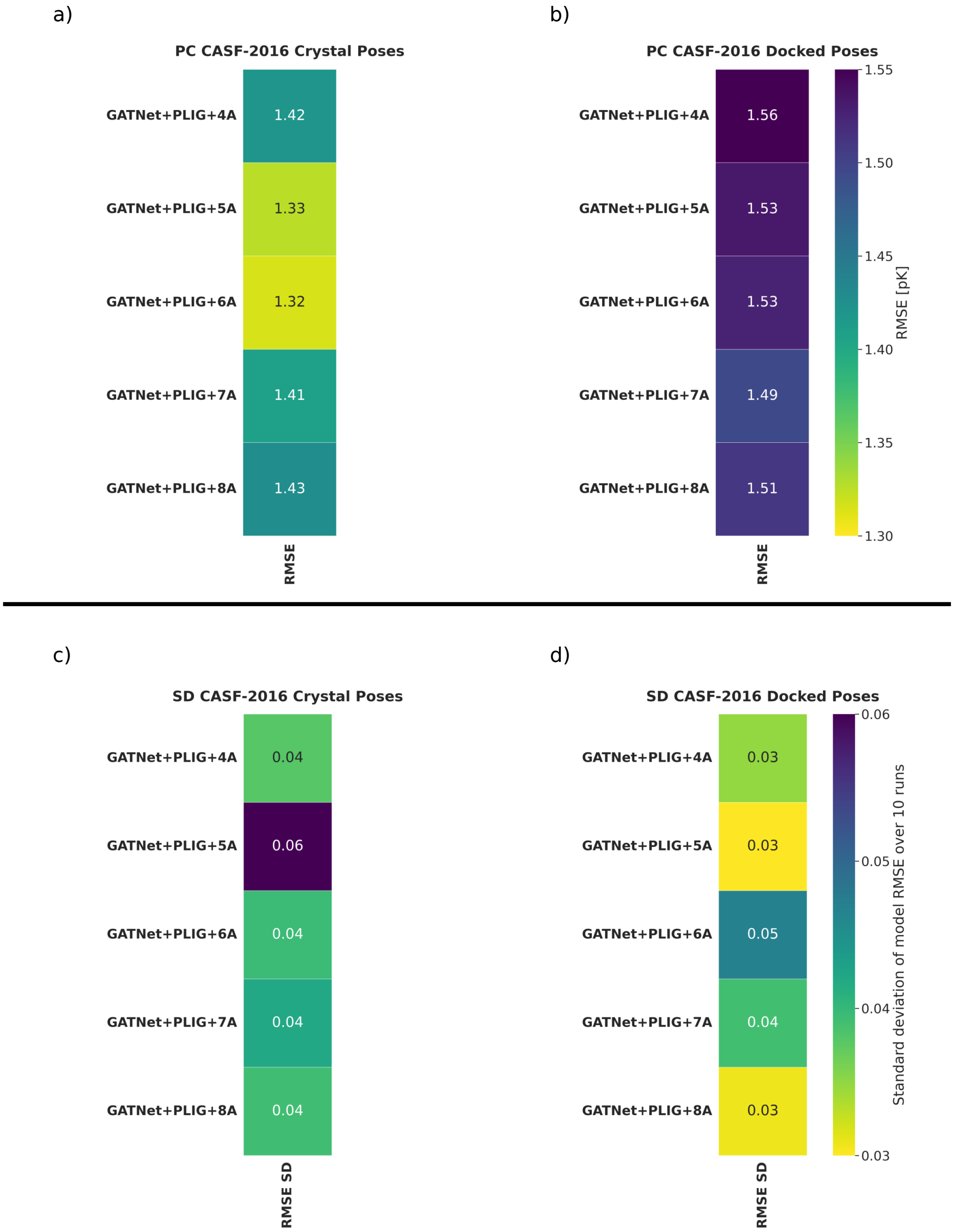

**Figure 5.4:** The reported root-mean-square error (RMSE) is calculated as the average over 10 train and test model iterations and shown in a-b with the corresponding standard deviation shown in c-d. All GATNet models were used without the second protein sequence embedding branch implementation. (a and c) RMSE and SD of the GATNet PLIG models with different proximity thresholds when trained and tested on crystal structures. (b and d) RMSE and SD of the GATNet PLIG models with different proximity thresholds when trained and tested on docked poses. Best performing threshold was the GATNet PLIG 5 and 6 Å models (RMSE = 1.33 and 1.32 for the 5 and 6 Å models, respectively). Model stability as measured by PC standard deviation varied between 0.03 (GATNet PLIG 4,5 and 8 Å trained/tested on docked poses) and 0.06 (GATNet PLIG 5 Å trained/tested on crystal poses).

a)

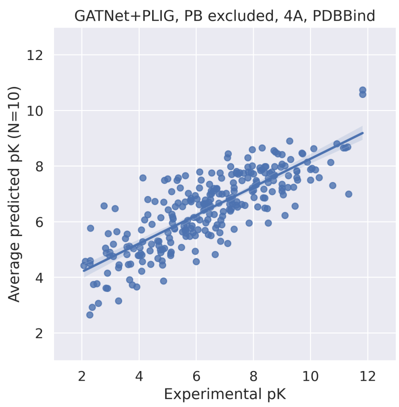

b)

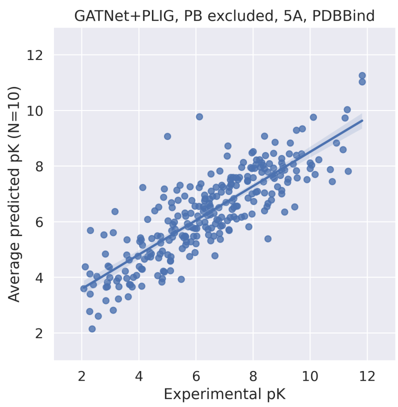

c)

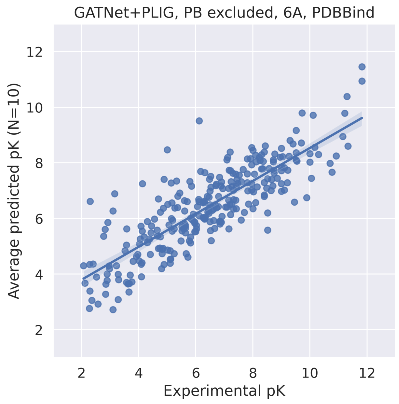

d)

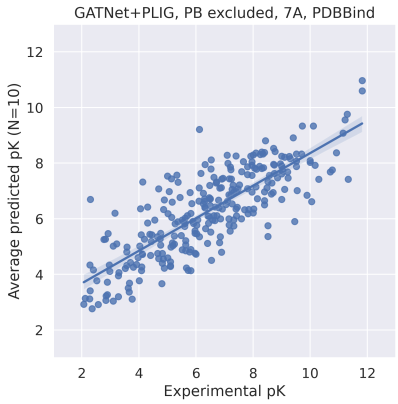

e)

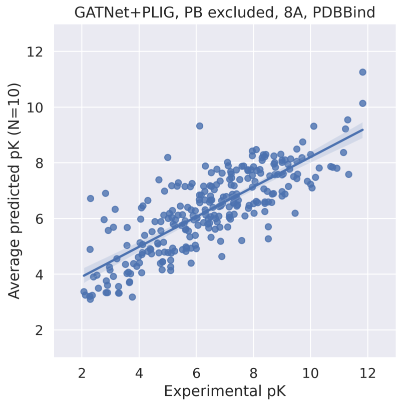

**Figure 5.5:** Scatter plots of the average prediction versus the experimentally determined pK value for each protein-ligand complex between the 10 model runs for the GATNet PLIG model at different proximity thresholds. All models were trained and tested on crystal structures. a) GATNet PLIG, no protein sequence, 4 Å threshold; b) GATNet PLIG, no protein sequence, 5 Å threshold; c) GATNet PLIG, no protein sequence, 6 Å threshold; d) GATNet PLIG, no protein sequence, 7 Å threshold; e) GATNet PLIG, no protein sequence, 8 Å threshold.

a)

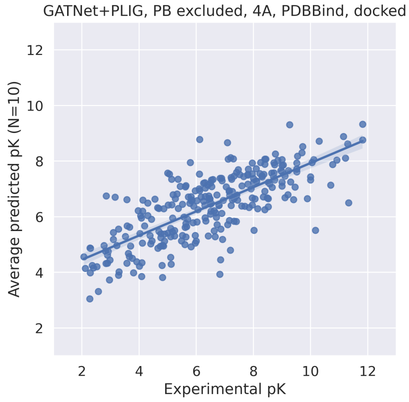

b)

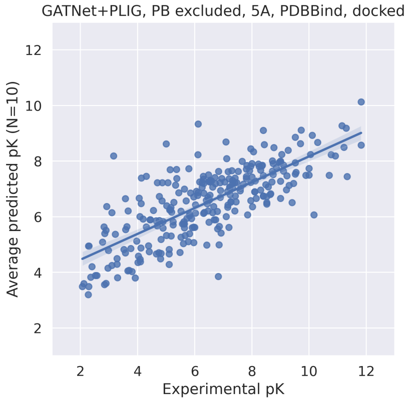

c)

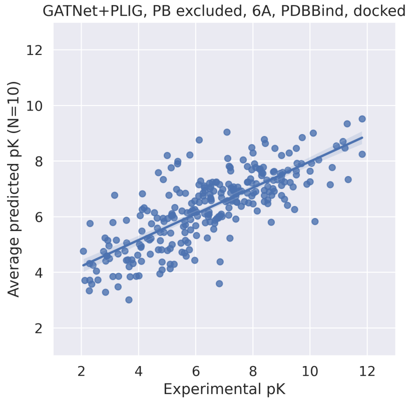

d)

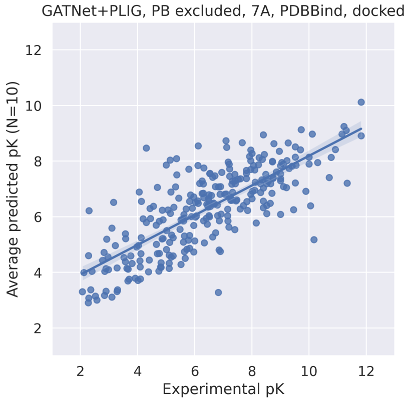

e)

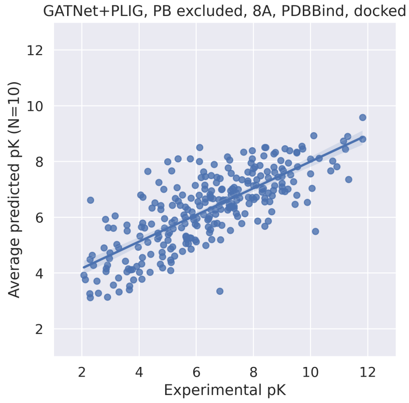

**Figure 5.6:** Scatter plots of the average prediction versus the experimentally determined pK value for each protein-ligand complex between the 10 model runs for the GATNet PLIG model at different proximity thresholds. All models were trained and tested on docked poses. a) GATNet PLIG, no protein sequence, 4 Å threshold; b) GATNet PLIG, no protein sequence, 5 Å threshold; c) GATNet PLIG, no protein sequence, 6 Å threshold; d) GATNet PLIG, no protein sequence, 7 Å threshold; e) GATNet PLIG, no protein sequence, 8 Å threshold.

### 6. *Docking results*

The quality of docked poses was estimated by calculating the symmetry-aware RMSD between all heavy atoms of the original crystal pose and the highest scoring docked pose for each protein-ligand complex using ODDT ([7]). A docked pose is considered to be of high quality if the calculated crystallographic RMSD is 2 Å or less. Overall, 39 % of docked poses were high quality. However, when splitting the dataset by the quality of the underlying crystal structures into the Refined Set and General Set, a subtle difference in pose quality is observed (42 % of docked poses from the Refined Set and 38 % of poses from the General Set, Figure 6.1). This might be due to the lower resolution of structures in the General Set where crystal structures can have resolutions of more than the 2 Å cutoff for pose quality estimation. In order to simulate a more realistic docking campaign where lower quality poses can be expected, we used all available docked poses as input for the scoring functions, regardless of pose accuracy.

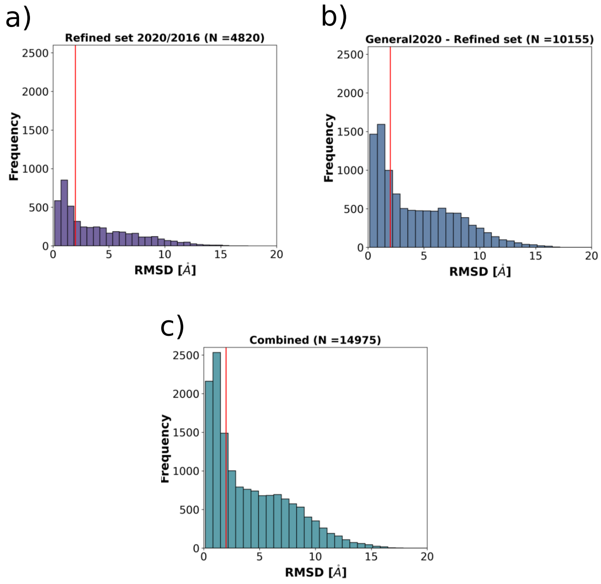

**Figure 6.1:** The red line marks the evaluation cutoff of 2 Å. All RMSD values were calculated as symmetry aware. A) Distribution of RMSD values (reported in Å) for the subset of the dataset that is part of the Refined Set 2020 and Refined Set 2016. B) Distribution of RMSD values (reported in Å) for the complexes in the General Set 2020 without complexes that are part of Refined Set 2020 or Refined Set 2016. C) Distribution of RMSD values (reported in Å) for the entire dataset (Refined Set 2016 and General Set 2020 including Refined Set 2020).

### 7. *Sequence Similarity Threshold Experiments*

In order to assess the ability of the best performing GATNet PLIG model to generalize between different protein families, protein-ligand pairs in the training dataset were eliminated based on their sequence identity to proteins represented in the CASF-2016 benchmark using 5 threshold levels between 50-100 % identity. The GATNet PLIG (no sequence) model was trained on the reduced dataset and tested against the full CASF-2016 dataset. The average PC and RMSE over 10 model runs as well as the corresponding standard deviation is shown in Figure 7.1. The scatter plots of the average prediction versus the experimentally determined pK value for each protein-ligand complex between the 10 model runs for each sequence similarity threshold are shown in Figure 7.2. The training dataset size for each threshold is shown in table 7.1. In addition to a decrease in performance with increasing strictness of the threshold, the standard deviation between model runs is increasing as well, indicating that models are more unstable as dataset size decreases and similarity between training and test set decreases.

| **Threshold** | **Training Set Size** |
|---|---:|
| Full training set | 14254 |
| 100 % threshold | 11917 |
| 95 % threshold | 11033 |
| 90 % threshold | 10969 |
| 70 % threshold | 10855 |
| 50 % threshold | 10408 |

**Table 7.1:** Size of the training set based on the sequence identity threshold.

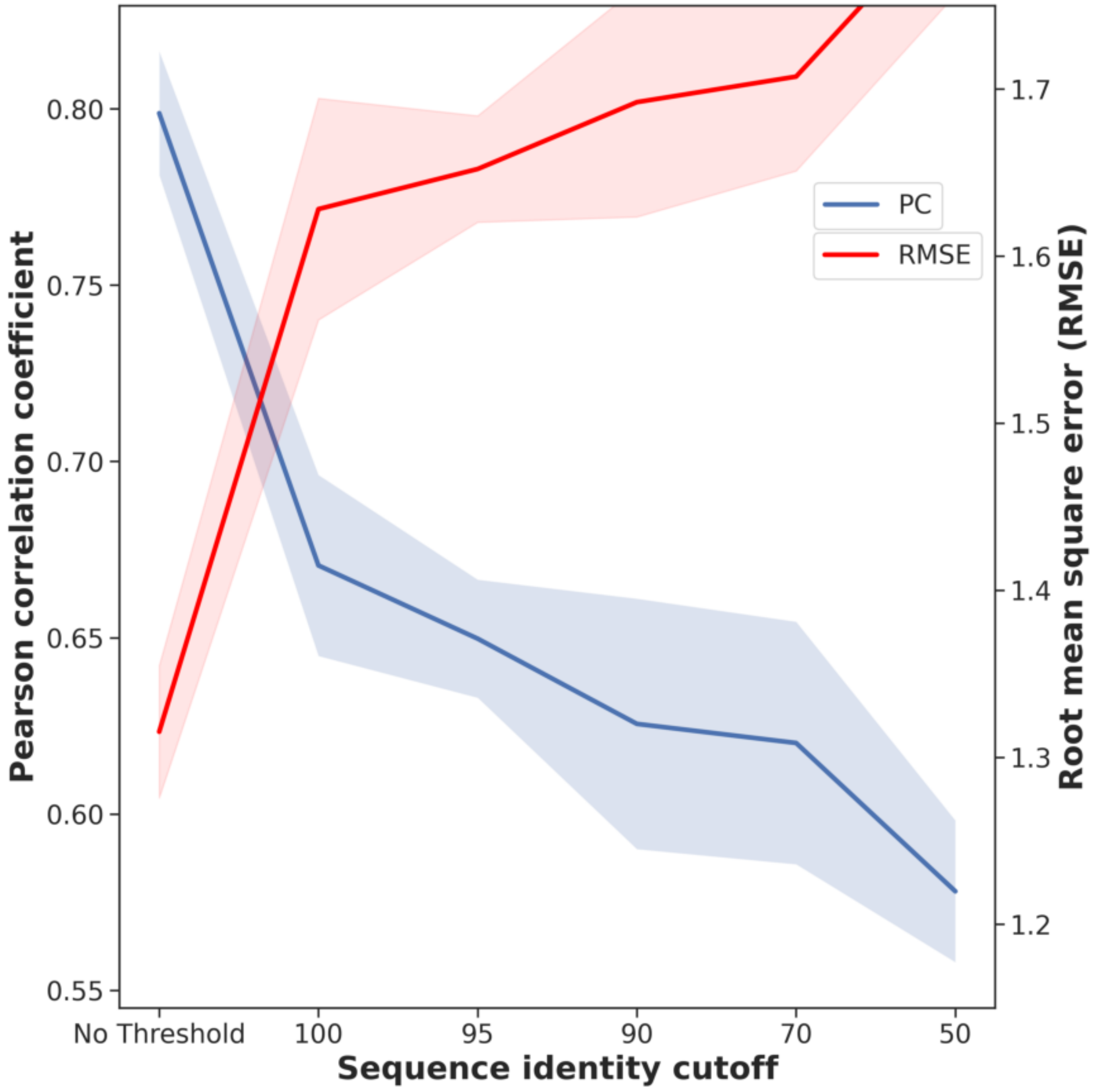

**Figure 7.1:** Pearson correlation coefficient ($\rho$) and Root-mean-square error (RMSE) of predicted versus experimental binding affinity for the GATNet PLIG model (no sequence, ensemble over 10 runs) when trained and tested on crystal poses. Protein-ligand complexes in the training set with a sequence identity at or above the cut-off value to proteins in the CASF-2016 test set were excluded resulting in a smaller dataset at every step.

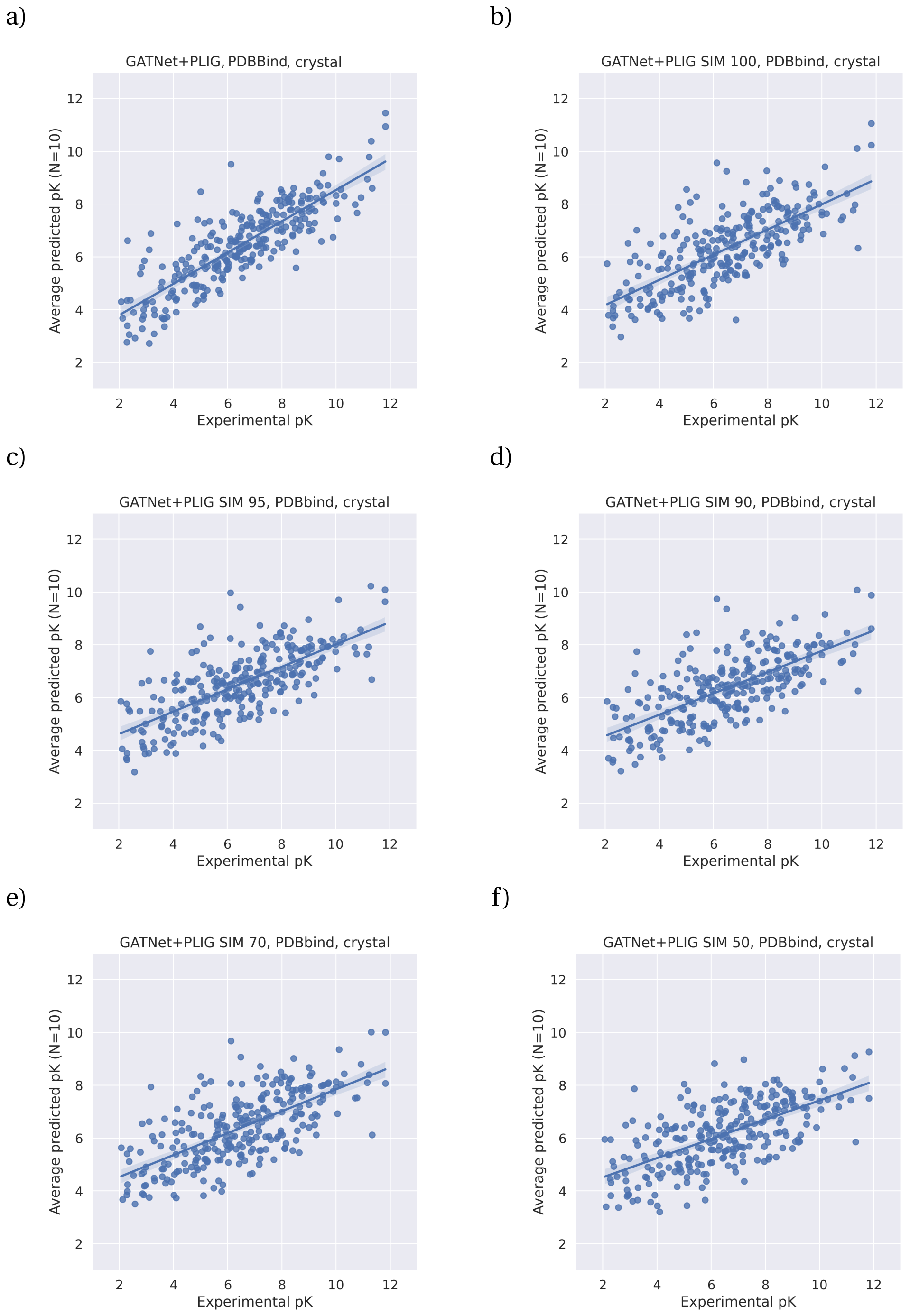

**Figure 7.2:** Scatter plots of the average prediction versus the experimentally determined pK value for each protein-ligand complex between the 10 model runs for each sequence similarity threshold. All models are GATNet PLIG models with no protein sequence embedding. a) No threshold, full training set; b) similarity threshold of 100%; c) similarity threshold of 95%; d) similarity threshold of 90%; e) similarity threshold of 70%; f) similarity threshold of 50%

### 8. *Bibliography*

[1] T. Akiba, S. Sano, T. Yanase, T. Ohta, and M. Koyama, “Optuna: A next-generation hyperparameter optimization framework,” in *Proceedings of the 25th ACM SIGKDD International Conference on Knowledge Discovery & Data Mining*, pp. 2623–2631, 2019.

[2] Z. Liu, Y. Li, L. Han, J. Li, J. Liu, Z. Zhao, W. Nie, Y. Liu, and R. Wang, “Pdb-wide collection of binding data: current status of the pdbbind database,” *Bioinformatics*, vol. 31, no. 3, pp. 405–412, 2015.

[3] Z. Liu, M. Su, L. Han, J. Liu, Q. Yang, Y. Li, and R. Wang, “Forging the basis for developing protein–ligand interaction scoring functions,” *Accounts of Chemical Research*, vol. 50, no. 2, pp. 302–309, 2017.

[4] F. Wu, A. Souza, T. Zhang, C. Fifty, T. Yu, and K. Weinberger, “Simplifying graph convolutional networks,” in *International Conference on Machine Learning*, 2019.

[5] J. Bergstra, R. Bardenet, Y. Bengio, and B. Kégl, “Algorithms for hyper-parameter optimization,” *Advances in Neural Information Processing Systems*, vol. 24, 2011.

[6] N. Sánchez-Cruz, J. L. Medina-Franco, J. Mestres, and X. Barril, “Extended connectivity interaction features: improving binding affinity prediction through chemical description,” *Bioinformatics*, vol. 37, pp. 1376–1382, 12 2020.

[7] M. Wójcikowski, P. Zielenkiewicz, and P. Siedlecki, “Open Drug Discovery Toolkit (ODDT): a new open-source player in the drug discovery field,” *Journal of Cheminformatics*, vol. 7, no. 1, p. 26, 2015.
